# Degeneracy in hippocampal physiology and plasticity

**DOI:** 10.1101/203943

**Authors:** Rahul Kumar Rathour, Rishikesh Narayanan

## Abstract

Degeneracy, defined as the ability of structurally disparate elements to perform analogous function, has largely been assessed from the perspective of maintaining robustness of physiology or plasticity. How does the framework of degeneracy assimilate into an encoding system where the ability to change is an essential ingredient for storing new incoming information? Could degeneracy maintain the balance between the apparently contradictory goals of the need to change for encoding and the need to resist change towards maintaining homeostasis? In this review, we explore these fundamental questions with the mammalian hippocampus as an example encoding system. We systematically catalog lines of evidence, spanning multiple scales of analysis, that demonstrate the expression of degeneracy in hippocampal physiology and plasticity. We assess the potential of degeneracy as a framework to achieve the conjoint goals of encoding and homeostasis without cross-interferences. We postulate that biological complexity, involving interactions among the numerous parameters spanning different scales of analysis, could establish disparate routes towards accomplishing these conjoint goals. These disparate routes then provide several degrees of freedom to the encoding-homeostasis system in accomplishing its tasks in an input- and state-dependent manner. Finally, the expression of degeneracy spanning multiple scales offers an ideal reconciliation to several outstanding controversies, through the recognition that the seemingly contradictory disparate observations are merely alternate routes that the system might recruit towards accomplishment of its goals. Against the backdrop of the ubiquitous prevalence of degeneracy and its strong links to evolution, it is perhaps apt to add a corollary to Theodosius Dobzhansky’s famous quote and state “nothing in physiology makes sense except in the light of degeneracy”.

**Highlights:**

- Degeneracy is the ability of structurally distinct elements to yield similar function
- We postulate a critical role for degeneracy in the emergence of stable encoding systems
- We catalog lines of evidence for the expression of degeneracy in the hippocampus
- We suggest avenues for research to explore degeneracy in stable encoding systems
- Dobzhansky wrote: “nothing in biology makes sense except in the light of evolution”
- A corollary: “nothing in physiology makes sense except in the light of degeneracy”

## 1. Introduction

The pervasive question on the relationship between structure and function spans every aspect of life, science and philosophy: from building architectures to the mind-body problem, from connectomics to genomics to proteomics, from subatomic structures to cosmic bodies and from biomechanics to climate science. Even within a limited perspective spanning only neuroscience, the question has been posed at every scale of brain organization spanning the genetic to behavioral ends of the spectrum. Efforts to address this question have resulted in extensive studies that have yielded insights about the critical roles of protein structure and localization, synaptic ultrastructure, dendritic morphology, microcircuit organization and large-scale synaptic connectivity in several neural and behavioral functions.

The question on the relationship between structure and function has spawned wide-ranging debates, with disparate approaches towards potential answers. At one extreme is the suggestion that structure defines function (Buzsaki, 2006):

> “The safest way to start speculating about the functions of a structure is to inspect its anatomical organization carefully. The dictum “structure defines function” never fails, although the architecture in itself is hardly ever sufficient to provide all the necessary clues.”

Within this framework, the following is considered as a route for understanding neural systems and behavior (Buzsaki, 2006):

> “First, we need to know the basic “design” of its circuitry at both microscopic and macroscopic levels. Second, we must decipher the rules governing interactions among neurons and neuronal systems that give rise to overt and covert behaviors.”

The other extreme is the assertion that “form follows function”, elucidated by Bert Sakmann (Sakmann, 2017), quoting Louis Sullivan:

> “Whether it be the sweeping eagle in his flight, or the open apple-blossom, the toiling work-horse, the blithe swan, the branching oak, the winding stream at its base, the drifting clouds, over all the coursing sun, form ever follows function, and this is the law. Where function does not change, form does not change”.

Within this framework, the approach to understanding neural structure function relations was elucidated as (Sakmann, 2017):

> “The approach we took, in order to discover structure-function relations that help to unravel simple design principles of cortical networks was, to first determine functions and then reconstruct the underlying morphology assuming that “form follows function”, a dictum of Louis Sullivan and also a Bauhaus design principle.”

A third approach embarks on addressing the structure-function question by recognizing the existence of ubiquitous variability and combinatorial complexity in biological systems. This was elucidated in a landmark review by Edelman and Gally, who presented an approach to structure-function relationship by defining degeneracy (Edelman and Gally, 2001):

> “Degeneracy is the ability of elements that are structurally different to perform the same function or yield the same output. Unlike redundancy, which occurs when the same function is performed by identical elements, degeneracy, which involves structurally different elements, may yield the same or different functions depending on the context in which it is expressed. It is a prominent property of gene networks, neural networks, and evolution itself. Indeed, there is mounting evidence that degeneracy is a ubiquitous property of biological systems at all levels of organization.”

They approach degeneracy and the structure-function question from an evolutionary perspective, noting (Edelman and Gally, 2001):

> “Here, we point out that degeneracy is a ubiquitous biological property and argue that it is a feature of complexity at genetic, cellular, system, and population levels. Furthermore, it is both necessary for, and an inevitable outcome of, natural selection.”

From this perspective, the supposition that a one-to-one relationship between structure and function exists is eliminated, thereby yielding more structural routes to achieving the same function. This perspective posits that biological complexity should be viewed from the evolutionarily advantageous perspective of providing functional robustness through degeneracy. Further, the degeneracy framework provides the system with higher degrees of freedom to recruit a state-dependent solution from a large repertoire of routes that are available to achieve the same function.

The advantages of biological variability (Foster *et al*., 1993; Gjorgjieva *et al*., 2016; Goldman *et al*., 2001; Katz, 2016; Marder, 2011; Marder and Goaillard, 2006; Marder *et al*., 2015; Marder and Taylor, 2011; O’Leary and Marder, 2014; Prinz *et al*., 2004; Taylor *et al*., 2009), degeneracy (Drion *et al*., 2015; Edelman and Gally, 2001; Leonardo, 2005; O’Leary *et al*., 2013; Whitacre and Bender, 2010; Whitacre, 2010) and complexity (Carlson and Doyle, 2002; Edelman and Gally, 2001; Stelling *et al*., 2004; Tononi *et al*., 1996, 1999; Weng *et al*., 1999; Whitacre, 2010), especially in terms of their roles in achieving robust function, have been widely studied and recognized in several biological process, including those in simple nervous systems. However, this recognition has been very limited in the mammalian neuroscience literature, where the focus is predominantly on explicitly assigning (or implicitly assuming) unique causal mechanistic relationships between constituent components and emergent functions. Here, we focus on the mammalian hippocampus, a brain region that has been implicated in spatial cognition, learning and memory, and review several lines of evidence that point to the existence of degeneracy in hippocampal physiology and plasticity. We argue that the elucidation of degeneracy spanning multiple scales could result in resolution of several existing controversies in the field, and provide an ideal setup to design experiments to understand neuronal systems, their adaptability and their responses to pathological insults.

The rest of the review is organized into four sections. In the first of these sections, we explore the foundations of degeneracy, especially from a perspective of an encoding system such as the hippocampus, and outline distinctions between different forms of homeostasis and their interactions with encoding-induced adaptations. In the second section, we build an argument that theoretical and experimental literature, spanning multiple scales of analysis, presents abundant support for the prevalence of degeneracy in almost all aspects of hippocampal physiology and plasticity. The third section explores the important question on the feasibility of establishing one-to-one structure-function relationships in systems that exhibit degeneracy through complexity. The final section concludes the review by briefly summarizing the arguments and postulates presented here on degeneracy in encoding within the degeneracy framework.

## 2. Degeneracy: Foundations from the perspective of an encoding system

Akin to the much broader span of physics from the subatomic to the cosmic scales, and very similar to studies on other biological systems, neural systems are studied at multiple scales of analysis (Fig. 1A). Although understanding neural systems *within* each of these scales of analysis is critical and has its own right for existence, a predominant proportion of neuro-scientific research is expended on *cross-scale emergence* of function through *interactions* among constituent components. One set of studies focus on the emergence of functions in a specified scale of analysis as a consequence of interactions among components in the immediately lower scale of analysis. An elegant example to such analysis is on the emergence of neuronal action potentials (a cellular scale function) as a consequence of interactions (Hodgkin and Huxley, 1952) between sodium and delayed rectifier potassium channels (molecular scale components). Another set of studies focus on the relationships between function at a specified scale of analysis and components that are integral to a scale that is several levels apart. With specific reference to the hippocampus, assessing the molecular-or cellular-scale components (*e.g*., receptors, synapses) that are *causally* responsible for learning and memory (a behavioral scale function that is several scales apart from the molecular/cellular scales) forms an ideal example for studies that belong in this category (Bliss and Collingridge, 1993; Kandel *et al*., 2014; Martin *et al*., 2000; Mayford *et al*., 2012; Neves *et al*., 2008a).

**Figure 1.**
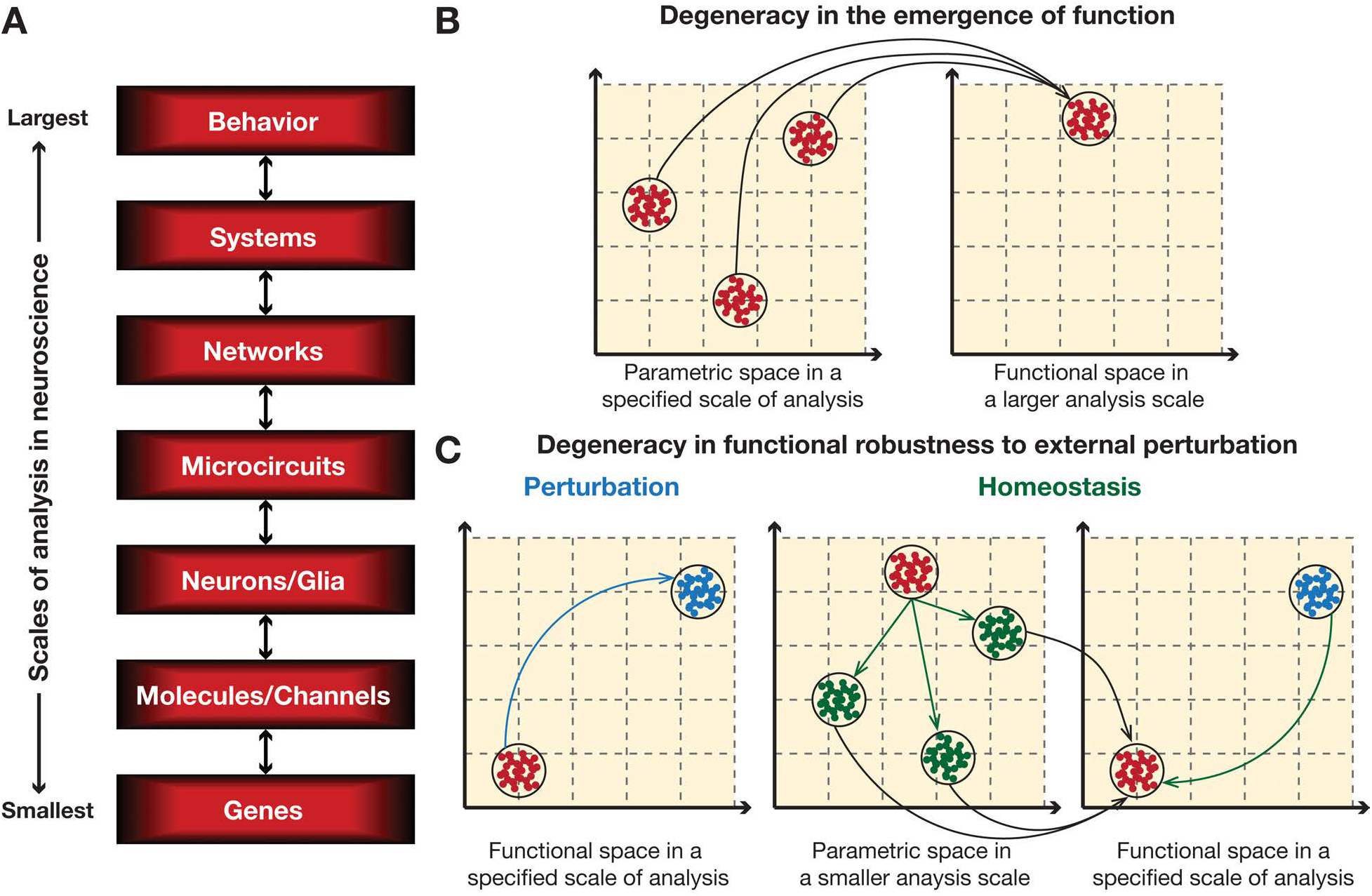
Degeneracy in the emergence of a function and its robustness to external perturbation across multiple scales of analysis. (A) Representation of multiple scales of analysis in neuroscience. The size (large and small) of the scale of analysis is representative of size of the constitutive components in that scale (Churchland and Sejnowski, 1992; Churchland and Sejnowski, 1988). (B) Disparate combinations of parameters in a specified scale of analysis could result in similar function in a larger scale of analysis. Each red circle in the smaller scale of analysis represents a combination of parameters that results in a specified function in large analysis scale, also represented by red circles there. The enclosing black circle in the larger scale represents experimentally observed variability in the function that is being assessed. On the other hand, the black circle in the smaller scale illustrates that robust functionality in the larger scale could be achieved even with small local perturbations in the parametric space. Larger perturbations beyond the black circle, however, would not yield robust functionality. The presence of multiple clusters of red circles in the smaller scale represents degeneracy, where similar functionality is achieved if parameters are within any of those multiple clusters. (C) Disparate combinations of parameters could compensate for functional impairment caused by external perturbation. *Left*, External perturbation results in the observed function in the larger scale of analysis switching from the baseline (red circles) to a perturbed state (blue circles). *Center*, In response, parameters in a smaller scale of analysis could undergo any of the several transitions, represented by green arrows, towards achieving functional homeostasis. Red circles represents the valid baseline parameters before perturbation, and green circles represent the state after the homeostatic response. *Right*, As a consequence of this homeostatic response involving any of the several disparate combinations of parameters, the system returns back to its baseline functionality (red circles).

Healthy and invigorating debates related to the philosophical and the scientific basis of such analyses, with themes ranging from broad discussions on reductionism vs. holism (Bennett and Hacker, 2003; Bickle, 2015; Jazayeri and Afraz, 2017; Krakauer *et al*., 2017; Panzeri *et al*., 2017) to more focused debates on the specific cellular components that are involved in specific aspects of coding and behavior (Bliss and Collingridge, 1993; Gallistel, 2017; Kandel *et al*., 2014; Kim and Linden, 2007; Martin *et al*., 2000; Mayford *et al*., 2012; Mozzachiodi and Byrne, 2010; Neves *et al*., 2008a; Otchy *et al*., 2015; Titley *et al*., 2017; Zhang and Linden, 2003), have contributed to our emerging understanding of neural systems and their links to behavior. Several studies have covered the breadth and depth of these debates (Bargmann and Marder, 2013; Bennett and Hacker, 2003; Bickle, 2015; Jazayeri and Afraz, 2017; Jonas and Kording, 2017; Kandel *et al*., 2014; Katz, 2016; Kim and Linden, 2007; Krakauer *et al*., 2017; Lazebnik, 2002; Marder, 1998, 2011, 2012; Marder *et al*., 2014; Marder and Thirumalai, 2002; Mayford *et al*., 2012; Panzeri *et al*., 2017; Tytell *et al*., 2011), and will not be the focus of this review.

Within the purview of degeneracy, the emergence of specific combinations of higher-scale functions (within the limits of biological variability) could be achieved (Fig. 1B) through interactions among disparate parametric combinations in a lower scale (Edelman and Gally, 2001; Foster *et al*., 1993; Gjorgjieva *et al*., 2016; Goldman *et al*., 2001; Marder, 2011; Marder and Goaillard, 2006; Marder *et al*., 2015; Marder and Taylor, 2011; O’Leary and Marder, 2014; Prinz *et al*., 2004; Rathour and Narayanan, 2012a, 2014; Srikanth and Narayanan, 2015; Stelling *et al*., 2004; Taylor *et al*., 2009). A straightforward corollary to this is that robust homeostasis in the maintenance of specific combinations of higher-scale functions in the face of perturbations there would be achieved through very different routes involving disparate parametric combinations in a lower scale (Fig. 1C). For instance, a change in neuronal firing rate at the cellular scale owing to external perturbations involving pathological insults or behavioral experience could be compensated for by different sets of changes to synaptic or intrinsic parameters (at the molecular scale) to achieve activity homeostasis (Gjorgjieva *et al*., 2016; Hengen *et al*., 2016; Nelson and Turrigiano, 2008; Turrigiano, 2011; Turrigiano, 1999, 2008; Turrigiano and Nelson, 2004). Thus, under the degeneracy framework, different uncorrelated clusters in the lower-scale parametric space could result in similar, if not identical, functional outcomes in the higher-scale measurement space, thereby suggesting a many-to-one relationship between the lower-scale parameters and higher-scale measurements (Edelman and Gally, 2001; Jazayeri and Afraz, 2017; Krakauer *et al*., 2017). Prominent lines of experimental evidence in support of degeneracy in neural systems have come from demonstrations of remarkable animal-to-animal variability in constituent components in providing analogous functional outcomes, and/or from results on many-to-many mappings between neural activity and behavior (Marder, 2011; Marder and Goaillard, 2006; Marder and Taylor, 2011; O’Leary and Marder, 2014; Schulz *et al*., 2006; Schulz *et al*., 2007; Vogelstein *et al*., 2014).

### 2.1. Degeneracy *vs*. compensation

A common misconception relating to degeneracy is that systems exhibiting degeneracy should compensate for the removal of a specific lower-scale component by recruiting other structural components there to yield the same higher-scale function. A corollary to this misconception is that an inability to compensate for the removal of a component is interpreted as evidence for the absence of degeneracy. For instance, consider an experiment where the “usefulness” of a specific gene is being tested by assessing deficits in a specific behavior after knockout of the gene under consideration. If the knockout resulted in the behavioral deficit, degeneracy is determined to be absent and the gene considered essential. On the other hand, for the case where there was no behavioral deficit, the gene is either considered non-essential or the result is interpreted as the expression of degeneracy where other components have compensated for the knockout.

There have been several warnings against such oversimplified interpretations, especially considering that biological systems are dynamic adaptive systems and not static (Edelman and Gally, 2001; Grashow *et al*., 2010; Marder, 2011; Marder and Goaillard, 2006; Marder and Taylor, 2011; O’Leary *et al*., 2014; Taylor *et al*., 2009; Wagner, 2005). Specifically, although the biological system adapts to the “unplanned” absence of the single gene (Edelman and Gally, 2001), it is not always essential that the adaptations result in compensation of one specific behavioral readout (of the several possible readouts (Jazayeri and Afraz, 2017; Krakauer *et al*., 2017)). Any compensation has been argued as a statistical result of the tradeoffs that are inherent to this complex, adaptive and nonlinear system that manifests degeneracy that is *emergent* across multiple scales of organization (Edelman and Gally, 2001; O’Leary *et al*., 2014). It has also been postulated that the compensatory process, and not the deletion, could have resulted in a specific deficit (O’Leary *et al*., 2014), especially because of the remarkable dissociation between different forms of homeostasis (see Sec. 2.2).

Further, especially given the ubiquitous variability across animals in terms of constituent components that elicit analogous behavior, it is clear that the impact of deletion of one specific component would be differential. This implies that the simplistic generalizability on the presence or absence of degeneracy based on a single parameter and a single measurement is untenable in complex adaptive systems. Additionally, with reference to the specific example of gene deletion, it is also important to distinguish between the acute impact of a lack of a protein that is tied to the gene and the developmental knockout (and associated compensatory mechanisms) of the specified gene (Edelman and Gally, 2001; Grashow *et al*., 2010; Marder, 2011; Marder and Goaillard, 2006; Marder and Taylor, 2011; O’Leary *et al*., 2014; Taylor *et al*., 2009).

In addition to these strong arguments against a one-to-one link between compensation and degeneracy, it is also important to consider the specifics of the expectations on the specific function that degeneracy is defined for and what functional deficit is to be compensated. Let’s consider the example of the emergence of membrane potential resonance in neurons as an example to illustrate this argument (Fig. 2). The emergence of resonance requires the expression of a resonating conductance, and the biophysical constraints on what makes a resonating conductance are well established (Cole, 1968; Das *et al*., 2017; Hodgkin and Huxley, 1952; Hutcheon and Yarom, 2000; Mauro, 1961; Mauro *et al*., 1970; Narayanan and Johnston, 2008). Hippocampal pyramidal neurons express several resonating conductances: the hyperpolarization-activated cyclic nucleotide-gated (HCN) nonspecific cation channels, the M-type potassium (KM) channels and the *T*-type calcium (CaT) channels, of which HCN and CaT channels exhibit overlapping voltage dependencies (Das *et al*., 2017; Hu *et al*., 2009; Hu *et al*., 2002; Narayanan and Johnston, 2007, 2008; Pike *et al*., 2000; Rathour and Narayanan, 2012a).

**Figure 2.**
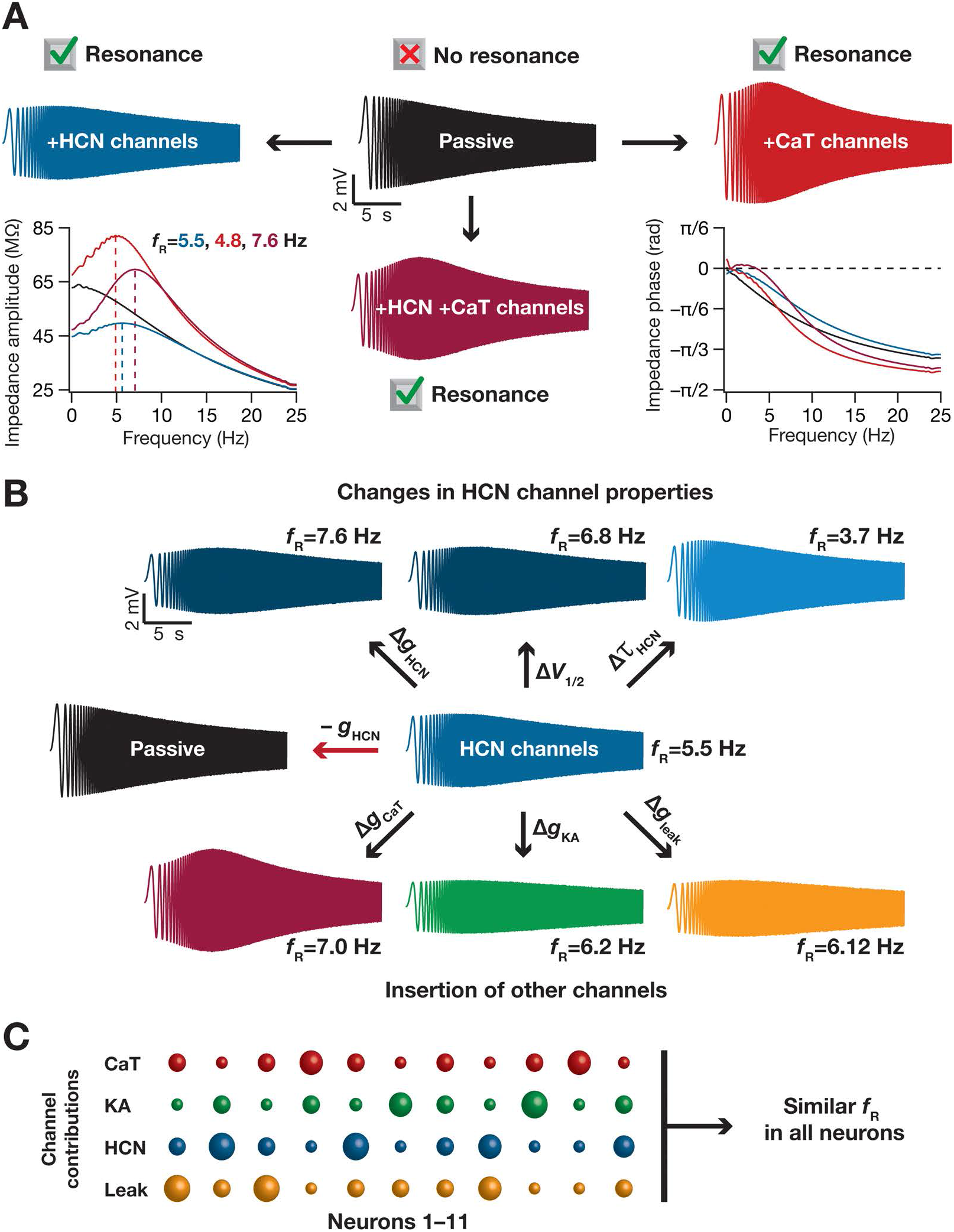
Qualitative vs. quantitative degeneracy. (A) Qualitative degeneracy, where the functional goal on which degeneracy is assessed is the expression of resonance, which could be achieved by the presence of one or more resonating conductances. Depicted are voltage traces obtained in response to a chirp current injection into neurons containing none, one or two resonating conductances. The hyperpolarization activated cyclic-nucleotide gated (HCN) and *T*-type calcium (CaT) are employed as the two example resonating conductances. In a neuron that expresses two or more resonating conductances (at sufficient densities), resonance ceases to express only when both resonating conductances are eliminated. The impedance amplitude (left bottom) and phase profiles (right bottom) are also shown for each color-matched chirp response. It may be noted that resonance in the amplitude profile and lead in the phase profile are observed when resonating conductances are expressed individually or together, and synergistically interact when they are expressed together. (B) Quantitative degeneracy, where the functional goal on which degeneracy is assessed is the ability to specify a target value of resonance frequency in the neuron, when a resonating conductance is expressed. Shown are some examples of the disparate possible routes to achieve quantitative changes to resonance frequency. One set of possibilities involves altering the properties of the channel mediating resonance (taken to be HCN in this example) such as its density (Δ*g*_HCN_), its gating properties (*e.g*., half-maximal activation voltage, Δ*V*_1/2_) or its kinetics (*e.g*., activation time constant, Δ*τ*_HCN_). The other set involves introducing *(e.g*., T-type calcium channels, Δ*g*_CaT_ or *A*-type potassium channels, Δ*g*_KA_) or altering (e.g., change in leak channels Δ*g*_leak_) other channels that modulate the resonance mediated by the resonating conductance (whose removal would abolish resonance, –*g*_HCN_, unless compensated by the expression of another resonating conductance). (C) In different neurons, the contribution of different channels to any measurement (shown here is resonance frequency, *f*_R_) could be variable. The size of each sphere scales with the quantum of contribution of a given channel (one among HCN, CaT, KA and leak) to *f*_R_ in a given neuron (11 neurons are depicted). Traces presented here and associated conclusions are drawn from previous studies (Hutcheon and Yarom, 2000; Narayanan and Johnston, 2007, 2008; Rathour *et al*., 2016; Rathour and Narayanan, 2012a).

Let’s first consider an example where the function on which degeneracy is assessed is qualitatively defined as the *expression* of membrane potential resonance (Fig. 2A). Whereas a passive neuron does not express resonance, the presence of the HCN and/or the CaT channels would result in the expression of resonance. This implies degeneracy in the function, where similar functionality (in this case, the expression of resonance) is through disparate components (channel combinations). In this scenario, depending on the variable expression profiles of HCN, CaT and other modulating channels, removal of only one of them could still result in the expression of resonance in specific neurons (Das *et al*., 2017; Rathour *et al*., 2016; Rathour and Narayanan, 2012a, 2014). However, removal of both HCN and CaT channels would result in a deficit in the assessed function, where resonance ceases to express. In this scenario, the requirement or usefulness of HCN or CaT channels to the expression of resonance is easily discernable by acute blockade experiments, although it would be difficult to predict (a) synergy between different channels that are expressed towards the emergence of resonance with such one-channel-at-a-time pharmacological blockade experiments; and (b) possible compensatory mechanisms involving changes in kinetics or voltage-dependence properties of other channels, say KM channels, in a double knockout scenario (Marder, 2011; Marder and Goaillard, 2006; O’Leary *et al*., 2014; Rathour and Narayanan, 2012a, 2014; Taylor *et al*., 2009).

In most encoding or homeostatic scenarios involving changes in constituent components, however, the functional outcome that is expected is a more quantitative readout of, say, firing rate or calcium concentration altered or returned to *specific values*. Therefore, a widely employed alternate interpretation (Foster *et al*., 1993; Goldman *et al*., 2001; Marder, 2011; Marder and Goaillard, 2006; Marder *et al*., 2015; Marder and Taylor, 2011; Prinz *et al*., 2004; Rathour and Narayanan, 2012a, 2014; Srikanth and Narayanan, 2015; Taylor *et al*., 2009) is where degeneracy is assessed as the ability of different structural components to elicit *quantitatively* similar functional measurements. With reference to our chosen example, this would translate to assessing degeneracy as the ability to achieve a *specific range of values* of resonance frequency with disparate combinations of parameters (Fig. 2B). If achieving a specific range of resonance frequency was the functional goal, and not the qualitative expression of resonance, then the possibilities are numerous. A resonating conductance is indeed required for the expression of resonance (Fig. 2B), but the goal is not to understand the expression of resonance, but to maintain resonance frequency at a specific value. In the presence of a resonating conductance, this goal could be achieved through very different structural routes either by altering other channel conductances or by altering properties of the resonating conductance itself. This implies the expression of degeneracy, where disparate parametric combinations could yield *quantitatively* similar resonance frequencies (Rathour and Narayanan, 2012a, 2014) across different models (Fig. 2C). Importantly, the order of degeneracy is rather large with the several active and passive properties, with the conductances, the voltage-dependence and kinetic properties of each of the several channels included. This also provides several routes to the emergence of compensation, where different channels and different parameters could differentially contribute to the emergence of similar functional measurements (Fig. 2C). We argue that this *quantitative* scenario with a large order of degeneracy is closer to the requirements of a system (at any given scale of organization) from the perspective of equilibrium and sustenance. The relevance of the qualitative scenario is rather limited to experiments that probe the expression of a specific phenomenon, which are “unplanned” from the evolutionary perspective there (Edelman and Gally, 2001).

Together, the question on the link between degeneracy and compensation should not be treated with simplistic ideas of linear interactions across components in a non-adapting system. The analyses should account for the specific definition of the function under consideration and the question on how degeneracy is defined. In addition, the nonlinear and synergistic interactions between different components that result in the specific function and animal-to-animal variability in expression profiles of constituent components should be assessed as part of such analyses. Finally, the possibility that “stochastic” compensatory process could be homeostatic or pathological and importantly on whether the challenge that is being posed to the system by the experiment is “planned” from the perspective of evolutionary convergence should also be considered (Edelman and Gally, 2001; Grashow *et al*., 2010; Marder, 2011; Marder and Taylor, 2011; O’Leary *et al*., 2014; Taylor *et al*., 2009).

### 2.2. Dissociation between different forms of homeostasis

It is clear from the examples presented above that the specific functional readout for which robustness or homeostasis ought to be maintained is a very critical question within the framework of degeneracy. Although degeneracy can be defined or observed with reference to any function at any scale of organization, the answer to the question on what specific functional homeostasis is absolutely essential from an evolutionary/neuroethological perspective isn’t clear. Even with reference to individual neurons, the literature has defined several forms of homeostasis (Gjorgjieva *et al*., 2016; Nelson and Turrigiano, 2008; Turrigiano, 2011; Turrigiano, 2008; Turrigiano and Nelson, 2004), with popular measures involving neuronal firing rate (Hengen *et al*., 2016), cytosolic calcium (Honnuraiah and Narayanan, 2013; O’Leary *et al*., 2014; Siegel *et al*., 1994; Srikanth and Narayanan, 2015) or excitation-inhibition balance (Yizhar *et al*., 2011). In addition, despite perpetual changes in afferent activity under *in vivo* conditions (Buzsaki, 2002, 2006, 2015; Srikanth and Narayanan, 2015; Tononi and Cirelli, 2006), specific neuronal subtypes maintain distinct functional signatures, say in terms of their excitability or oscillatory or frequency selectivity measurements, that are different from other neuronal subtypes even in the same brain region (Hoffman *et al*., 1997; Migliore and Shepherd, 2002, 2005; Narayanan and Johnston, 2007, 2008; Pike *et al*., 2000; Spruston, 2008; Zemankovics *et al*., 2010). Further, synaptic properties such as strength and release probabilities are also very discernable across different synaptic subtypes (say excitatory *vs*. inhibitory) even on the same postsynaptic neuron (Andrasfalvy and Magee, 2001; Andrasfalvy and Mody, 2006; Dittman *et al*., 2000; Koester and Johnston, 2005; Magee and Cook, 2000; Smith *et al*., 2003). This suggests the existence of some form of homeostasis that maintains these intrinsic and synaptic measurements, including or apart from firing rate or calcium homeostasis or excitatory-inhibitory balance, despite behaviorally driven encoding changes or perpetual activity switches that are common in the hippocampus and other regions of the brain. Does maintenance of one of them translate to maintenance of all of them? If not, which of these different forms of homeostasis are absolutely essential for the animal from the evolutionary/neuroethological perspective?

There are several lines of clear evidence that there are remarkable dissociations between different forms of homeostasis (Srikanth and Narayanan, 2015). First, cellular-or network-scale functions could robustly emerge with disparate combinations of molecular-or cellular-scale parameters (Foster *et al*., 1993; Marder, 2011; Marder and Goaillard, 2006; Prinz *et al*., 2004; Rathour and Narayanan, 2014; Taylor *et al*., 2009). These observations suggest that precise homeostatic balance at a lower scale (e.g., ion channels expressed to exact conductance values) is not essential for maintaining functional homeostasis at a higher scale. Second, even in the same set of neurons/networks/animals, different measurements have different dependencies on underlying parameters, and these dependencies could be variable. For instance, in the same neuron, resonance frequency could have a larger dependence on one channel subtype with input resistance being critically regulated by another channel, with the specifics of these dependencies variable across different neurons of the same subtype (Fig. 3A). Studies have shown that different channels could have differential and variable impact on disparate measurements from the same neuron, even in a location dependent manner (Grashow *et al*., 2010; O’Leary *et al*., 2014; Rathour and Narayanan, 2014; Taylor *et al*., 2009). Additionally, acute blockade of one specific channel results in weakly correlated changes in different measurements in the same neuron (Rathour *et al*., 2016). This implies that changing individual constitutive components to maintain robust homeostasis in one of the measurements does not necessarily translate to robust homeostasis in all the other measurements.

**Figure 3.**
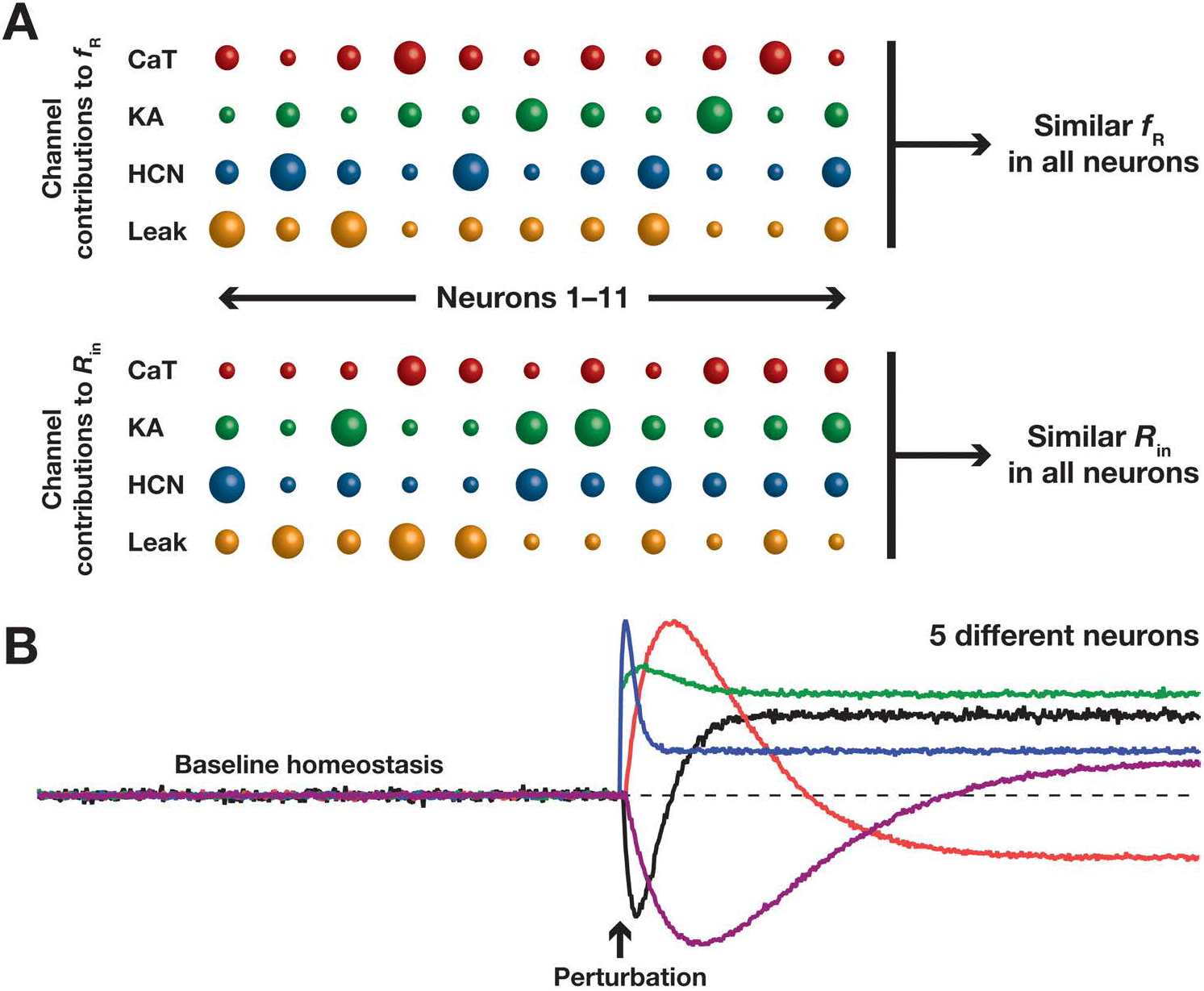
Dissociation between different forms of homeostasis. (A) In different neurons, the contribution of different channels to different measurements (shown here are resonance frequency,fR, and input resistance, *R*_in_) is differential and variable. The size of each sphere scales with the quantum of contribution of a given channel (one among HCN, CaT, KA and leak) to *f*_R_ in a given neuron (11 neurons are depicted). It may be noted that in any given neuron, it is not necessary that the contributions of any given channel to *f*_R_ and *R*_in_ need not be equal, even when both *f*_R_ and *R*_in_ are similar across all neurons. Cartoon illustrations are derived from data presented in previous studies (Rathour *et al*., 2016; Rathour and Narayanan, 2012a, 2014; Srikanth and Narayanan, 2015). (B) Although baseline homeostasis is efficaciously maintained in five different neurons, their responses to an identical perturbation need not necessarily be identical or even similar. The perturbation could be a plasticity-inducing stimulus driven by behavioral experience or by pathological conditions. Cartoon illustration was derived from analyses presented in previous studies (Anirudhan and Narayanan, 2015; O’Leary *et al*., 2014; Srikanth and Narayanan, 2015).

Third, for maintenance of calcium homeostasis across neurons in a network or in neurons that are subjected to perpetual switches in afferent activity, it is not essential that functional homeostasis across different intrinsic or synaptic measurements is maintained. Specifically, owing to inherent variability in different constitutive components, the channel conductance values or neuronal intrinsic properties or synaptic strengths could be very different across different neurons despite maintenance of precise calcium homeostasis in neurons or their network (Gjorgjieva *et al*., 2016; O’Leary *et al*., 2014; Srikanth and Narayanan, 2015). Finally, calcium and firing rate homeostasis have been shown to be dissociated whereby tremendous variability in channel conductance values, firing rate and pattern of firing have been observed despite efficacious maintenance of calcium homeostasis (O’Leary *et al*., 2013; O’Leary *et al*., 2014; Srikanth and Narayanan, 2015). Together, these studies establish that none of the individual forms of homeostasis (in calcium concentration or in channel densities channel or in intrinsic functional characteristics including neuronal firing-rate) necessarily translate to or follow from any other among them (O’Leary *et al*., 2013; O’Leary *et al*., 2014; Rathour and Narayanan, 2012a, 2014; Srikanth and Narayanan, 2015), implying clear dissociations between different forms of homeostasis.

### 2.3. Baseline *vs*. plasticity profile homeostasis

An important and necessary cynosure in the physiology of encoding systems is their ability to change in a manner that promotes adaptability to the environment. In other words, the ability to undergo plasticity is an important requirement for it to encode or learn newly available information from the environment. Such plasticity has been shown to be ubiquitous, spanning cellular and network structures across almost all regions, and could be triggered by development (Desai *et al*., 2002; Desai *et al*., 1999; Luo and Flanagan, 2007; Schreiner and Winer, 2007; Turrigiano and Nelson, 2004; White and Fitzpatrick, 2007), by learning processes (Kandel, 2001; Kandel *et al*., 2014; Kim and Linden, 2007; Lamprecht and LeDoux, 2004; Narayanan and Johnston, 2012; Titley *et al*., 2017; Zhang and Linden, 2003) or by pathological insults (Beck and Yaari, 2008; Bernard *et al*., 2007; Brager and Johnston, 2014; Grant, 2012; Johnston *et al*., 2016; Kullmann, 2002; Lee and Jan, 2012; Lehmann-Horn and Jurkat-Rott, 1999; Lerche *et al*., 2013; Poolos and Johnston, 2012). A traditional method to study such plasticity mechanisms is to subject neuronal or synaptic structures to specific activity patterns towards understanding the rules for plasticity in specific components. Assessed through such protocols, distinct synapses show signature profiles of plasticity in terms of the strength and direction of synaptic plasticity elicited by specific activity patterns. Additionally, there are also specific sets of non-synaptic forms of plasticity (in channel densities and properties, for instance) that are concomitant to the synaptic plasticity induced by different activity patterns (Abbott and Nelson, 2000; Abbott and Regehr, 2004; Bi and Poo, 1998; Bliss and Collingridge, 1993; Bliss and Lomo, 1973; Chung *et al*., 2009a; Chung *et al*., 2009b; Cooper and Bear, 2012; Dittman *et al*., 2000; Dudek and Bear, 1992; Fortune and Rose, 2001; Frick *et al*., 2004; Jorntell and Hansel, 2006; Lin *et al*., 2008; Losonczy *et al*., 2008; Lujan *et al*., 2009; Magee and Johnston, 1997; Markram *et al*., 1997; Narayanan and Johnston, 2007, 2008; Shah *et al*., 2010; Sjostrom *et al*., 2008). This implies *plasticity profile homeostasis* (Anirudhan and Narayanan, 2015; Mukunda and Narayanan, 2017), where synapses of the same subtype respond similarly to analogous afferent activity, thereby resulting in a subtype-dependent rule for synaptic plasticity (Larsen and Sjostrom, 2015). In terms of non-synaptic plasticity, such plasticity profile homeostasis could be generalized to subtypes of cells manifesting specific forms of neuronal plasticity (in intrinsic properties, for instance).

Juxtaposed against the considerable variability in different constitutive components across neurons of the same subtype, and given the critical dissociations between different forms of homeostasis (Sec. 2.2), it is easy to deduce that the maintenance of baseline homeostasis of a given measurement (say activity or calcium) does not necessarily imply that the system will respond in a similar manner to identical perturbations (Fig. 3B). As the direction and strength of change in activity or calcium is a critical determinant of the plasticity profile (Lisman, 1989; Lisman *et al*., 2002; Lisman *et al*., 2012; Lisman, 2001; Nevian and Sakmann, 2006; Regehr, 2012; Shouval *et al*., 2002; Sjostrom and Nelson, 2002; Sjostrom *et al*., 2008; Zucker, 1999; Zucker and Regehr, 2002), variable responses to incoming perturbations (physiological or pathophysiological) would translate to very distinct plasticity profiles even in synapses of the same subtype (Anirudhan and Narayanan, 2015; Mukunda and Narayanan, 2017; O’Leary *et al*., 2013; Srikanth and Narayanan, 2015). Therefore, from the perspective of homeostasis in encoding systems such as the hippocampus, it is not just sufficient to ask if baseline homeostasis of a given measurement is maintained. It is also important to ask if the response of the system to identical perturbations is similar to enable plasticity profile homeostasis. The absence of such plasticity profile homeostasis would result in very different adaptations to identical perturbations even under baseline conditions, resulting in the absence of signature plasticity profiles being associated with specific neurons and synapses. Although there is dissociation between the maintenance of baseline *vs*. plasticity profile homeostasis, studies have demonstrated degeneracy in the maintenance of short- and long-term plasticity profiles. Specifically, these studies have shown that disparate combinations of ion channel conductances and calcium-handling mechanisms could yield analogous short-or long-term plasticity profiles (Anirudhan and Narayanan, 2015; Mukunda and Narayanan, 2017). Although we dealt with plasticity profile homeostasis and its dissociation from baseline homeostasis, a related phenomenon that involves plasticity of *plasticity profiles* has been defined as metaplasticity (Abraham, 2008; Abraham and Bear, 1996; Abraham and Tate, 1997; Cooper and Bear, 2012; Hulme *et al*., 2013; Sehgal *et al*., 2013). Lines of evidence supporting degeneracy in hippocampal metaplasticity and its roles in stable learning will be explored in Sec. 3.3.

### 2.4. Encoding and homeostasis within the degeneracy framework

The function of learning systems extends beyond simple maintenance of physiological or plasticity homeostasis. The functional goal in these systems is rather contrary to *maintenance* of homeostasis, because encoding or learning of new information demands *alteration* in physiology/behavior through continual *adaptation* in an experience-/activity-dependent manner. This presents a paradoxical requirement where components ought to *change* to encode new information, *without* perturbing the overall homeostatic balance of the system. Thus, encoding of a new experience entails a tricky balance between change and homeostasis (James, 1890):

> “Plasticity, then, in the wide sense of the word, means the possession of a structure weak enough to yield to an influence, but strong enough not to yield all at once. Each relatively stable phase of equilibrium in such a structure is marked by what we may call a new set of habits.”

From the degeneracy and physiology perspectives, this balance poses several tricky questions that the literature does not present definitive answers to. For instance, could learning systems accomplish this balance between encoding of new information *and* maintenance of homeostasis *within* the framework of degeneracy? In other words, could the plasticity mechanisms that define encoding *and* the homeostatic mechanisms that negate the impact of perturbation *together* be realized through disparate combinations of constitutive components (Narayanan and Johnston, 2012; Nelson and Turrigiano, 2008; Turrigiano, 2007, 2011; Turrigiano *et al*., 1994; Turrigiano, 1999; Turrigiano and Nelson, 2000)? Would the availability of more routes to achieve encoding or homeostasis be detrimental or be advantageous towards accomplishing these goals together? Would the dissociations between different forms of homeostasis (Sec. 2.2) and between baseline vs. plasticity profile homeostasis (Sec. 2.3) translate to severe constraints on accomplishing this balance *within* the framework of degeneracy?

Together, there are lines of evidence supporting the formulation that plasticity and homeostasis individually could be achieved through several non-unique routes through disparate combinations of constituent components (Anirudhan and Narayanan, 2015; Mukunda and Narayanan, 2017; Narayanan and Johnston, 2012; Nelson and Turrigiano, 2008; O’Leary *et al*., 2013; Srikanth and Narayanan, 2015; Turrigiano, 2007, 2011; Turrigiano *et al*., 1994; Turrigiano, 1999; Turrigiano and Nelson, 2000). However, the focus on achieving the *conjoined* goals of effectuating changes in response to new information *and* maintaining robust homeostasis in the face of such changes *within* the framework of degeneracy have been conspicuously lacking. Such focus is especially important because of the seemingly contradictory requirements of the two processes, where one necessitates change and the other works to negate any change, resulting in the possibility where there could be detrimental cross-interference working towards negating each other. Therefore, for the framework of degeneracy to be relevant in learning systems, it is important that future studies assess the twin goals of encoding and homeostasis to be synergistically conjoined rather than treat them as isolated processes that independently achieve their respective goals. Without the recognition of such synergy between encoding and homeostatic systems, assessing the ability of these two processes to avoid cross-interference becomes intractable.

### 2.5. Curse-of-dimensionality or evolutionary robustness

Curse of dimensionality, coined by Bellman (Bellman, 1957), refers to the extreme difficulties encountered with the comprehension or solution to a problem that involves exorbitantly large numbers of input variables, their attributes and possible solutions. In biology in general, and in neuroscience in particular, the dimensions of the parametric space is typically large, making dimensions of the interactional space (the space that covers all forms of interactions spanning all these parameters) even larger. The variability of parametric values even in systems exhibiting similar functions and the perpetual adaptation of these parameters in response to external perturbations (or even baseline turnover towards maintaining homeostasis) make it impossible to localize any biological function to a small subspace of this large interactional space. This, as a consequence of the curse of dimensionality, translates to mathematical and computational intractability of biological systems because of insufficiency of collected data towards providing an accurate answer to questions related to comprehending or assessing the system.

The framework of degeneracy on the other hand suggests that biological systems thrive on this parametric and interactional complexity because it provides the ideal substrate for arriving at disparate structural routes to robust functional similarity. Several strong qualitative and quantitative arguments, based on several lines of evidence spanning different scales of analysis across different biological systems, have been placed in favor of synergistic links between degeneracy, complexity, robustness, evolvability and adaptation. Therefore, the dimensionality of the parametric and interactional space of biological systems should not be treated as a curse in terms of our inability to analytically track or comprehend the system, but as a fundamental and necessary feature towards achieving the contradictory yet conjoint goals (Sec. 2.4) of functional robustness (Edelman and Gally, 2001; Kitano, 2007; Marder, 2011; Marder and Goaillard, 2006; Rathour *et al*., 2016; Rathour and Narayanan, 2012a, 2014; Sporns *et al*., 2000; Stelling *et al*., 2004; Tononi and Cirelli, 2006; Tononi and Edelman, 1998; Tononi *et al*., 1998; Wagner, 2005, 2008), evolvability (Edelman and Gally, 2001; Wagner, 2008; Whitacre and Bender, 2010; Whitacre, 2010) and adaptation (Albantakis *et al*., 2014; Anirudhan and Narayanan, 2015; Joshi *et al*., 2013; Mukunda and Narayanan, 2017).

Importantly, the recognition of the critical links between complexity, degeneracy and adaptability allows for better design of experimental and analysis techniques for assessing biological systems and their function. Not only do these techniques alleviate the pains of hand tuning in computational models (Prinz *et al*., 2003), but also recognize the implications for parametric variability to robust functions and the fallacies associated with misinterpretation of results from knockout animals in the face of perpetual biological compensation (Edelman and Gally, 2001; Grashow *et al*., 2010; Marder, 2011; Marder and Goaillard, 2006; Marder and Taylor, 2011; O’Leary *et al*., 2014; Taylor *et al*., 2009; Wagner, 2005). Some classes of techniques developed with the recognition of the strong links between variability, complexity, adaptability, degeneracy and robustness are: (a) the global sensitivity analysis technique (Sec. 3.2) that employs a stochastic search algorithm spanning a large parametric space and optimizes for *multiple* physiological objectives (Foster *et al*., 1993; Goldman *et al*., 2001; Marder, 2011; Marder and Goaillard, 2006; Marder and Taylor, 2011; Prinz *et al*., 2004; Rathour and Narayanan, 2014); (b) the theoretical and experimental assessment of the links between quantitative complexity measures and robustness with reference to several physiological and pathophysiological attributes (Albantakis *et al*., 2014; Edelman and Gally, 2001; Joshi *et al*., 2013; Kitano, 2007; Sarasso *et al*., 2015; Sporns *et al*., 2000; Stelling *et al*., 2004; Tononi and Edelman, 1998; Tononi *et al*., 1998; Tononi *et al*., 1996, 1999; Wagner, 2005, 2008; Whitacre and Bender, 2010; Whitacre, 2010); and (c) plasticity models that have accounted for concomitant changes in multiple components (Secs. 3.6–3.7) rather than focusing on a one-to-one relationship between functional plasticity and one specific component that undergoes changes (Abbott and LeMasson, 1993; Anirudhan and Narayanan, 2015; LeMasson *et al*., 1993; Mukunda and Narayanan, 2017; O’Leary *et al*., 2013; O’Leary *et al*., 2014; Siegel *et al*., 1994; Srikanth and Narayanan, 2015). These analyses have made it abundantly clear that the complexities inherent to biological systems should be considered as substrates for functional robustness through degeneracy (Edelman and Gally, 2001), rather than be viewed from the curse-of-dimensionality perspective.

### 2.6. Error correction mechanisms

A critical requirement in a system that is endowed with degeneracy is an error-correcting feedback mechanism that regulates constituent components in an effort to achieve a specific function. For instance, consider the example where the goal is to achieve calcium homeostasis in a neuron. In this scenario, as the specific regulatory mechanism that is to be triggered is dependent on the current state of the neuron, or more precisely the current levels of calcium, it is important that the regulatory mechanism is geared towards *correcting* the *error* between the target function and the current state (Abbott and LeMasson, 1993; LeMasson *et al*., 1993; O’Leary *et al*., 2013; O’Leary *et al*., 2014; Siegel *et al*., 1994; Srikanth and Narayanan, 2015). This requires a closed circuit feedback loop that initiates a compensatory mechanism that is driven by the quantitative distance between the target function and the current state. This state-dependent perpetual error correction becomes especially important in a scenario where distinct regulatory mechanisms govern the different constitute components. With the specific example at hand, let’s say the error correcting feedback mechanism regulates ion channel conductances by altering their protein expression through several transcription factors (Srikanth and Narayanan, 2015). In such a scenario, calcium homeostasis could be achieved by recruiting several non-unique sets of these transcription factors. As each of these transcription factors could be coupled to the regulation of distinct combinations of ion channels, calcium homeostasis could be achieved through several non-unique combinations of ion channels.

Within the degeneracy framework, although distinct solutions are possible with weak pairwise correlations between constitutive components, there is a strong synergistic *collective* dependence of these components to achieve a function (Rathour and Narayanan, 2014). Specifically, let’s consider two neurons (neurons 1 and 2) with distinct sets of non-unique parametric combinations that yielded very similar function. However, given the nonlinearities of neural systems, it would be infeasible to expect similar function from a third neuron built with one-half of the parameters taken from neuron 1 and the other half taken from neuron 2. This collective cross-dependence is an essential component of systems manifesting degeneracy and should be respected by mechanisms that regulate the constitutive components. Returning to specific example under consideration, the specific *ensemble* of the targeted transcription factors and channel conductances are important in terms of which solution is *chosen* within the degeneracy framework. This places strong requirements on the distinct regulatory mechanisms, transcription factors in this case, that they strongly interact with each other rather than acting independent of each other (Srikanth and Narayanan, 2015) in a manner that is *driven* by the error that is being fed back in a state-dependent temporally precise manner.

These requirements become especially important in an encoding system such as the hippocampus, whose afferent activity is perpetually variable in a behavioral state-dependent manner, requiring temporally proximal feedback for the continuous maintenance of robust function. A simple solution to account for cross-interacting regulatory mechanisms is to assume the existence of only one regulatory mechanism that governs all constitutive components (e.g., one transcription factors controls all channels and receptors on a neuron (O’Leary *et al*., 2014)). However, this might not always be valid or possible or feasible (Srikanth and Narayanan, 2015), especially if the complexity of system is enormous (e.g., coexistence of multiple transcription factors in the hippocampus (Alberini, 2009; Bading *et al*., 1993; Dolmetsch, 2003; Lein *et al*., 2007). In these scenarios, it is important that the error-sensing and regulatory mechanisms also exhibit degeneracy and are strongly inter-coupled to each other through cross-regulatory mechanisms at that scale as well (*e.g*., multiple calcium sensors accompanied by a network of transcription factors coupled through feedback loops that regulate each other (Cheong *et al*., 2011; Kotaleski and Blackwell, 2010; Losick and Desplan, 2008; Thattai and van Oudenaarden, 2001; Yu *et al*., 2008)). In summary, the ability to achieve functional robustness through degeneracy in any scale of analysis requires continuous correction of functional deficits, without which it is impossible to adjudge the efficacious accomplishment of a desired goal through a chosen route (which is one among the many possible routes). In a system with enormous complexity, this is typically achieved through an error-correcting feedback pathway that recruits multiple cross-interacting regulatory mechanisms towards maintaining collective cross dependence of constituent mechanisms (Rathour and Narayanan, 2014; Srikanth and Narayanan, 2015).

## 3. Degeneracy at multiple scales in the hippocampus

The hippocampus is a brain region that has been shown to be critically involved in spatial representation of the external environment and in several forms of learning and memory (Anderson *et al*., 2007; Eichenbaum, 2012; Hartley *et al*., 2014; Moser *et al*., 2008; Neves *et al*., 2008a; Scoville and Milner, 1957). As a region that is involved in encoding of new information and one that is part of the medial temporal lobe that is critically sensitive to excitotoxic insults (Bernard *et al*., 2007; Dam, 1980; de Lanerolle *et al*., 1989; Johnston *et al*., 2016; Sloviter, 1991), it is important that the hippocampal cells maintain some form of activity homeostasis to avoid runaway excitation.

The hippocampus consists of several subtypes of neurons and glia receiving afferent information from tens of thousands of synapses and expressing distinct sets of a wide variety of ligand-gated receptors and voltage-gated ion channels, each built through complex structural interactions between a number of main and auxiliary subunits (Lai and Jan, 2006; Migliore and Shepherd, 2002; Nusser, 2009, 2012; Vacher *et al*., 2008; Verkhratsky and Steinhauser, 2000). The regulatory role of glial cells and their constitutive components in synaptic information processing is well established (Allen and Barres, 2005, 2009; Araque, 2008; Araque *et al*., 2014; Araque *et al*., 1999; Bazargani and Attwell, 2016; Deitmer *et al*., 2006; Fields and Stevens-Graham, 2002; Halassa *et al*., 2007; Halassa and Haydon, 2010; Haydon and Carmignoto, 2006; Pannasch and Rouach, 2013; Pascual *et al*., 2005; Perea and Araque, 2005; Perea *et al*., 2009), providing additional structural substrates that could participate in the encoding and homeostasis processes. The basic properties and regulation of these and other membrane and cytoplasmic protein structures, in conjunction with intracellular (including the ER and the trafficking apparatus) and intercellular interaction dynamics (including neuronal synaptic connectivity and the glial syncytium) and morphological characteristics, regulates the intricate balance between encoding and homeostasis within the hippocampal structure. In addition to these, hippocampal structure and function are critically reliant on the afferent and efferent connectivity patterns, the metabolic pathways that drive and interact with the local cellular structures and the several forms of state-dependent modifications to each of these components. Together, the combinatorial complexity of the constitutive components that define hippocampal function is staggeringly astronomical.

A fundamental question that is of considerable interest to the research community is on how the hippocampus achieves robust function, especially in accomplishing the apparently contradictory goals of adaptive change and homeostasis (Sec. 2), in the face of such combinatorial complexity that drives its physiology and plasticity. Within the framework of degeneracy, it could be argued that the complexity is an enabler, and not an impediment, towards achieving functional robustness.

Does hippocampal physiology manifest degeneracy at multiple scales, whereby similar hippocampal function could be achieved through disparate structural combinations? In this section, we view hippocampal research spanning the past several decades through the lens of degeneracy and present clear qualitative and quantitative lines of evidence arguing for the ubiquitous presence of degeneracy spanning multiple scales of hippocampal function. We review lines of evidence showing multiple routes to achieving several critical hippocampal functions, which in some cases have been considered to be lines of evidence that are in apparent contradiction to each other, triggering expansive debates and arguments within the field. In a manner similar to (Edelman and Gally, 2001), we systematically explore the expression of degeneracy at distinct scales (starting at the molecular scale and moving incrementally to the systems/behavioral scale) of hippocampal function (Fig. 1A), with function(s) or physiological measurements assessed within the specified scale of analysis. We postulate that the recognition of the ubiquitous prevalence of degeneracy would provide an evolutionarily routed framework to unify the several apparently contradictory routes to achieving the same function as necessity, rather than luxury, towards achieving physiological robustness.

### 3.1. Degeneracy in the properties of channels and receptors

Hippocampal neurons are endowed with myriad voltage and ligand dependent ion channels, with well-defined gradients in their expression profiles and their properties (Barnard *et al*., 1998; Dingledine *et al*., 1999; Johnston and Narayanan, 2008; Magee and Cook, 2000; Migliore and Shepherd, 2002; Narayanan and Johnston, 2012; Paoletti *et al*., 2013; Sieghart and Sperk, 2002). The presence of these channels, with their signature characteristics and expression profiles, has been shown to play critical roles in the physiology (Das *et al*., 2017; Johnston *et al*., 1996; Johnston and Narayanan, 2008; Magee, 2000; Narayanan and Johnston, 2012), plasticity (Frick and Johnston, 2005; Johnston *et al*., 2003; Remy *et al*., 2010; Shah *et al*., 2010; Sjostrom *et al*., 2008) and pathophysiology (Bernard *et al*., 2007; Brager and Johnston, 2014; Johnston *et al*., 2016; Kullmann, 2002; Lee and Jan, 2012; Lerche *et al*., 2013) of hippocampal neurons and their networks. Therefore, it is essential that the biophysical properties and expression profiles of these channels be tightly regulated to ensure functional robustness.

The regulation of targeting, localization and properties of these channels at specific levels, however, is a problem that involves several degrees of combinatorial freedom. The reasons behind this complexity are manifold. First, most of these channels are not protein molecules derived from single genes, but are assembled from several possible pore-forming and auxiliary subunits, expressed in different stoichiometry (Catterall, 1993, 1995; Gurnett and Campbell, 1996; Hille, 2001; Isom *et al*., 1994). The presence or absence of a specific pore-forming or auxiliary subunit, and the specific ratios of their expression are important for trafficking, localization and properties of these channels. For instance, *A*-type K^+^ channels in the hippocampus could be assembled by the main subunits from the Kv1 or Kv4 families and auxiliary subunits from the KChIP and DPP families (Amarillo *et al*., 2008; Birnbaum *et al*., 2004; Jerng *et al*., 2004; Kim *et al*., 2007; Kim *et al*., 2005; Sun *et al*., 2011; Vacher and Trimmer, 2011), whereas auxiliary subunits MiRP1, KCR1 and TRIP8b have been implicated in regulating trafficking and properties of *h* channels assembled with main subunits from the HCN family of proteins. Additionally, the properties of *h* channels, in terms of their voltage-dependence, their kinetics and modulation by cyclic nucleotides, are critically regulated by the specific isoforms that are expressed in conjunction with the specific stoichiometry of such expression (Biel *et al*., 2009; He *et al*., 2014; Lewis *et al*., 2011; Much *et al*., 2003; Robinson and Siegelbaum, 2003; Santoro *et al*., 2000; Santoro *et al*., 2009; Santoro *et al*., 2004; Ulens and Siegelbaum, 2003; Ulens and Tytgat, 2001; Zolles *et al*., 2009).

Second, targeting and functional properties of these assembled channels (Trimmer and Rhodes, 2004; Vacher *et al*., 2008) could be critically modulated by different forms of post-translational modification (Derkach *et al*., 1999; Derkach *et al*., 2007; Levitan, 1994; Misonou *et al*., 2004; Much *et al*., 2003; Shah *et al*., 2010; Sjostrom *et al*., 2008), by local pH (Holzer, 2009), by interaction with intracellular messengers (Armstrong and Bezanilla, 1974) and by lipid composition of the plasma membrane (Levitan and Barrantes, 2012). For instance, trafficking of *A*-type K^+^ channels is phospho-regulated in a manner that is dependent on their main and auxiliary subunits (Birnbaum *et al*., 2004; Hammond *et al*., 2008; Lin *et al*., 2011; Lin *et al*., 2010; Vacher and Trimmer, 2011), and differences between proximal and distal dendritic sodium channels are partly mediated by phosphorylation states of these channels (Gasparini and Magee, 2002).

Third, distinct channels have been demonstrated to have structural interactions with each other, thereby cross-regulating the functional properties of each other. For instance, structural interactions between Cav3 and Kv4 channel families are known to regulate neuronal activity through efficient transfer of calcium influx from Cav3 channels to bind onto KChIPs that modulate Kv4 channel function (Anderson *et al*., 2010). Finally, these channels can undergo activity-dependent plasticity and neuromodulation (Biel *et al*., 2009; Cantrell and Catterall, 2001; He *et al*., 2014; Hoffman and Johnston, 1999; Lee and Dan, 2012; Marder, 2012; Marder *et al*., 2014; Marder and Thirumalai, 2002; Robinson and Siegelbaum, 2003), which also could result in important changes to their trafficking and functional properties (Sec. 3.6).

How do these channels maintain specific location-dependent levels of expression with specific properties despite this staggering complexity that results in their assemblage and specific function? From the description above, it is clear that channels achieve specific properties and localization through multiple structural routes involving several subunits, enzymes associated with post-translational modification, neuromodulators and their receptors and several signaling cascades (also see Sec. 3.4–3.6). This follows the observation that each functional property of the channel, including its localization and targeting, is regulated by multiple mechanisms, each endowed with the ability to bidirectionally modulate the functional property. Therefore, the combinatorial complexity of regulation and the involvement of different structural routes to achieve similar function together provide ample lines of evidence for the expression of degeneracy in achieving specific function for channels and receptors expressed in the hippocampus. In answering the question on how robustness might be achieved, the argument within the framework of degeneracy would be that functional robustness in the assemblage, targeting and function of ion channels is achieved as a *consequence* of the underlying regulatory and interactional complexity.

### 3.2. Degeneracy in neuronal physiological properties

The presence of various ligand and voltage dependent ion channels confers signature neurophysiological properties, such as input resistance, firing rate, frequency selectivity and integration and propagation of potentials across axonal and dendritic processes, upon different hippocampal neurons (Hutcheon and Yarom, 2000; Johnston *et al*., 1996; Llinas, 1988). Although there is remarkable variability in these measurements even within a single neuronal subtype (Dougherty *et al*., 2012; Dougherty *et al*., 2013; Malik *et al*., 2016), different neuronal subtypes within the same subregion have signature electrophysiological characteristics (Anderson P, 2007; Freund and Buzsaki, 1996; Klausberger and Somogyi, 2008; Spruston, 2008) that are maintained despite the combinatorial complexity of ion channels expressed in these neurons. Additionally, prominent relationships between intrinsic neurophysiological properties and various pathological conditions, including epilepsy and Fragile X mental disorder, have been reported across several neurological disorders (Beck and Yaari, 2008; Bernard *et al*., 2007; Brager and Johnston, 2014; Johnston *et al*., 2016; Kullmann, 2002; Lee and Jan, 2012; Lehmann-Horn and Jurkat-Rott, 1999; Lerche *et al*., 2013; Poolos and Johnston, 2012). Thus, from the maintaining robust physiology and from the perspective of avoiding pathological excitability conditions, it is essential that neurons maintain their signature electrophysiological characteristics.

It is now recognized across systems that there is no one-to-one relationship between neurophysiological properties and the channels that regulate them (Sec. 2.1–2.3, Fig. 2–3). It is established that several channels contribute to the emergence and regulation of a specific physiological property, and the same channel could regulate several physiological properties, resulting in a many-to-many mapping between channels and physiological properties. In addition to the example assessing degeneracy in resonance properties (Sec. 2.1–2.2, Fig. 2–3), we could also consider the example of maintaining neuronal firing rates at specific levels. Whereas fast Na^+^ and delayed rectifier K^+^ channels mediate action potential firing in hippocampal neurons, their firing rate profiles are regulated by an array of ion channels including the *A*-type K^+^, HCN, GIRK, *M*-type K^+^ and SK channels (Adelman *et al*., 2012; Gasparini and DiFrancesco, 1997; Gu *et al*., 2005; Hu *et al*., 2007; Kim and Johnston, 2015; Kim *et al*., 2005; Malik and Johnston, 2017; Narayanan and Johnston, 2007; Rathour *et al*., 2016).

These observations provide specific insights about the relationship between channels and physiological properties (Sec. 2.1–2.3; Fig. 2–3). First, there is degeneracy in the emergence of neurophysiological properties, where disparate combinations of channels could come together to elicit similar functional properties (Das *et al*., 2017; Drion *et al*., 2015; Foster *et al*., 1993; Goldman *et al*., 2001; Marder, 2011; Marder and Goaillard, 2006; Rathour *et al*., 2016; Rathour and Narayanan, 2012a, 2014; Taylor *et al*., 2009).

Second, the dependence of different physiological properties on distinct channels is variable even within the same neuronal subtype, and is a function of the variable expression profiles of these channels (Drion *et al*., 2015; O’Leary *et al*., 2014; Rathour and Narayanan, 2014; Taylor *et al*., 2009). For instance, whereas *A*-type K^+^ channels might contribute maximally to maintaining firing rates at a specific level in one neuron, in another neuron of the same subtype it could be SK channels.

Third, the dependence of different physiological properties in the *same* neuron on distinct channels is differential and variable, where pharmacological blockade of one channel may have a stronger effect on a specific physiological property compared to another (Rathour *et al*., 2016). As a consequence of these observations, there is a dissociation between robust maintenance of one physiological property and that of another (Srikanth and Narayanan, 2015). Maintenance of only a few physiological properties would not necessarily translate to maintenance of all physiologically relevant properties. All relevant physiological properties ought to be explicitly maintained for overall robustness.

Fourth, hippocampal neurons are endowed with complex dendritic arborization with several well-defined functional maps expressing along their somato-dendritic arbor, making proteostasis, or protein homeostasis (Balch *et al*., 2008), in these neurons a complex problem (Hanus and Schuman, 2013; Narayanan and Johnston, 2012). Despite the strong structural constraint of maintaining robustness of several tightly coupled location-dependent functional measurements, it has been demonstrated that it is not essential to maintain individual channels at specific densities or with specific properties for achieving robust functional homeostasis. Instead, several disparate combinations of channel parameters, spanning properties and densities of several channels, could robustly maintain *concomitant* homeostasis of multiple functions across the dendritic arbor (Rathour and Narayanan, 2014). It is however essential to note that dendritic morphology plays a crucial role in regulating intrinsic properties and their location-dependent characteristics, especially in electrotonically *non*-compact hippocampal pyramidal neurons (Dhupia *et al*., 2015; Golding *et al*., 2005; Krichmar *et al*., 2002; Mainen and Sejnowski, 1996; Narayanan and Chattarji, 2010; Spruston *et al*., 1994; Spruston *et al*., 1993), and could contribute to degeneracy in the emergence of single-neuron physiology.

Finally, depending on the localization profiles and voltage-dependent properties of different channels they may or may not spatiotemporally interact (Migliore and Migliore, 2012; Mishra and Narayanan, 2015; Rathour and Narayanan, 2012b). For instance, owing to mostly non-overlapping voltage-dependence and localization profiles, *M*-type K^+^ and HCN channels mediate complementary somato-dendritic theta filtering in hippocampal neurons (Hu *et al*., 2009; Narayanan and Johnston, 2007, 2008). In contrast, *A*-type K^+^ and HCN channels strongly overlap both in their voltage-dependence and localization, resulting in their ability to co-regulate the same form of resonance in hippocampal pyramidal neurons (Rathour *et al*., 2016; Rathour and Narayanan, 2012a, 2014)

These insights are driven by experimental observations coupled with physiologically relevant computational models that allowed greater flexibility in terms of understanding mechanistic basis, importance of ion channel interactions and the degree of contribution of each channel type in regulating neuronal properties. Multi parametric multi objective stochastic search algorithms are a class of algorithms that has been employed as an extremely effective method to explore cellular-level degeneracy in a systematic and rigorous manner through global sensitivity analysis (Anirudhan and Narayanan, 2015; Drion *et al*., 2015; Foster *et al*., 1993; Goldman *et al*., 2001; Mukunda and Narayanan, 2017; Rathour and Narayanan, 2012a, 2014; Taylor *et al*., 2009). These algorithms provide a quantitative route to understanding the structure of the global parametric space in any given model, without making explicit assumptions about co-variation of different parameters test the robustness of the system to parametric variability. In this technique, model neurons generated by uniform random sampling of the global parametric space are tested against experimental statistics of several measurements. Model neurons that satisfy *several* experimental constraints are declared as “valid models”. The use of multiple measurements to establish the validity of models is essential because of afore-mentioned (Sec. 2.1–2.3) dissociation between different forms of homeostasis and the differential dependence of different measurements on distinct constitutive components (Fig. 2–3). It is well recognized in the design principle of these techniques that establishing physiological equivalence of only a partial set of measurements *does not* necessarily ensure that the other measurements which have not been constrained by the validation process are within the physiological ranges (Achard and De Schutter, 2006; Foster *et al*., 1993; Goldman *et al*., 2001; Hobbs and Hooper, 2008; Marder, 2011; Marder and Goaillard, 2006; Marder and Taylor, 2011; Prinz *et al*., 2003; Prinz *et al*., 2004; Rathour and Narayanan, 2012a, 2014; Srikanth and Narayanan, 2015; Taylor *et al*., 2009; Tobin *et al*., 2006; Weaver and Wearne, 2008). If such a stochastic search algorithm fails to yield any valid model that satisfies all the physiological objectives, the interpretation should not be that the specified model configuration is incapable of achieving all objectives. This is because the stochastic search does not *entirely* span the global parametric space, thereby allowing for the possibility that valid solutions could exist within the unexamined regions of this parametric space.

Once the validity of a (typically small) subset of models through multiple physiological constraints is established, the approach has been employed to explore degeneracy by assessing pair-wise and cross-dependencies across different parameters. Pairwise correlations across valid model parametric values are typically employed to explore such dependencies, where a strong correlation between any two parameters is interpreted as a pointer to potential co-regulation of biological mechanisms defining these parameters (Anirudhan and Narayanan, 2015; Foster *et al*., 1993; Goldman *et al*., 2001; Mukunda and Narayanan, 2017; Rathour and Narayanan, 2012a, 2014; Taylor *et al*., 2009). These analyses also provide insights about how critically specific parameters should be regulated to achieve the *multiple* objectives imposed by the validation criteria. Importantly, these algorithms provide a quantitative route to finding the relative sensitivities of different measurements to each channel that contributed to the emergence of robust functionality spanning multiple measurements. It is recognized that the dependence of measurements on individual channels would be variable given that different model neurons are endowed with considerable variability in each channel conductance. However, it is still known that the average dependence of a given measurement (say resonance frequency) is higher for one specific channel (say HCN channels), *relative* to the other channels expressed in the system. Different methodologies have been proposed to assess these relative contributions and have been effectively employed to understand the differential and variable dependencies of different measurements on each underlying channel (O’Leary *et al*., 2014; Rathour and Narayanan, 2014; Taylor *et al*., 2009).

Together, through a confluence of electrophysiological and computational techniques that assessed variability and homeostasis in neuronal and channel properties, the expression of degeneracy in the emergence of single neuron physiology is well established across several systems, including the mammalian hippocampus. It is clear that disparate combinations of morphological and channel parameters could robustly yield analogous single neuron physiology, despite being constrained by *multiple* measurements that span the entire somato-dendritic arbor of the *same* neuron.

### 3.3. Degeneracy in calcium regulation and in the induction of synaptic plasticity

Whereas the ability to maintain baseline physiological measurements at specific levels is important from the homeostasis perspective, the ability to alter responses (through changes in parameters) towards achieving a specific target is important from the perspective of learning or encoding. This ability to undergo long-term plasticity is absolutely critical in an encoding system. One of the most well studied forms of long-term plasticity in hippocampal neurons is plasticity in synaptic structures. There are several lines of evidence for degeneracy in the induction, expression and maintenance of long-term synaptic plasticity and the mechanisms that are associated with each of these distinct phases of synaptic plasticity. As long-term synaptic plasticity is relatively well studied, we will first outline these lines of evidence from the synaptic plasticity perspective and then switch to the implications for *concomitant* non-synaptic plasticity that typically accompanies synaptic plasticity.

A popular methodology to study long-term synaptic plasticity in neurons within the hippocampus and other brain structures is the use of specific induction protocols that result in synaptic plasticity. These induction protocols are activity-dependent, and are typically induced by combinations of presynaptic stimulation and/or postsynaptic current injection. There are also several chemical protocols for inducing synaptic plasticity, say through depolarization induced through elevated levels of extracellular potassium or potassium channel blockers (Hanse and Gustafsson, 1994; Huang and Malenka, 1993; Huber *et al*., 1995; Lin *et al*., 2008; Otmakhov *et al*., 2004; Roth-Alpermann *et al*., 2006). These protocols are critically tied to the specific synaptic structures that are studied and show signature profiles across synaptic structures of similar subtypes (Abbott and Nelson, 2000). The protocols required for induction of synaptic plasticity are not unique. Several disparate protocols with very distinct combinations of presynaptic stimulation and/or postsynaptic current injection (Fig. 4) have been shown to elicit long-term potentiation (LTP) or long-term depression (LTD). The cellular mechanisms required for inducing LTP are also very different across these protocols, with differences sometimes manifesting even within a single protocol for synapses at two different locations on the same neuron. For instance, with the theta burst protocol for inducing LTP (Fig. 4A), proximal synaptic LTP requires pairing with backpropagating action potentials, but distal synapses recruit dendritic spikes and do not require backpropagating action potentials (Golding *et al*., 2002; Kim *et al*., 2015; Magee and Johnston, 1997).

**Figure 4.**
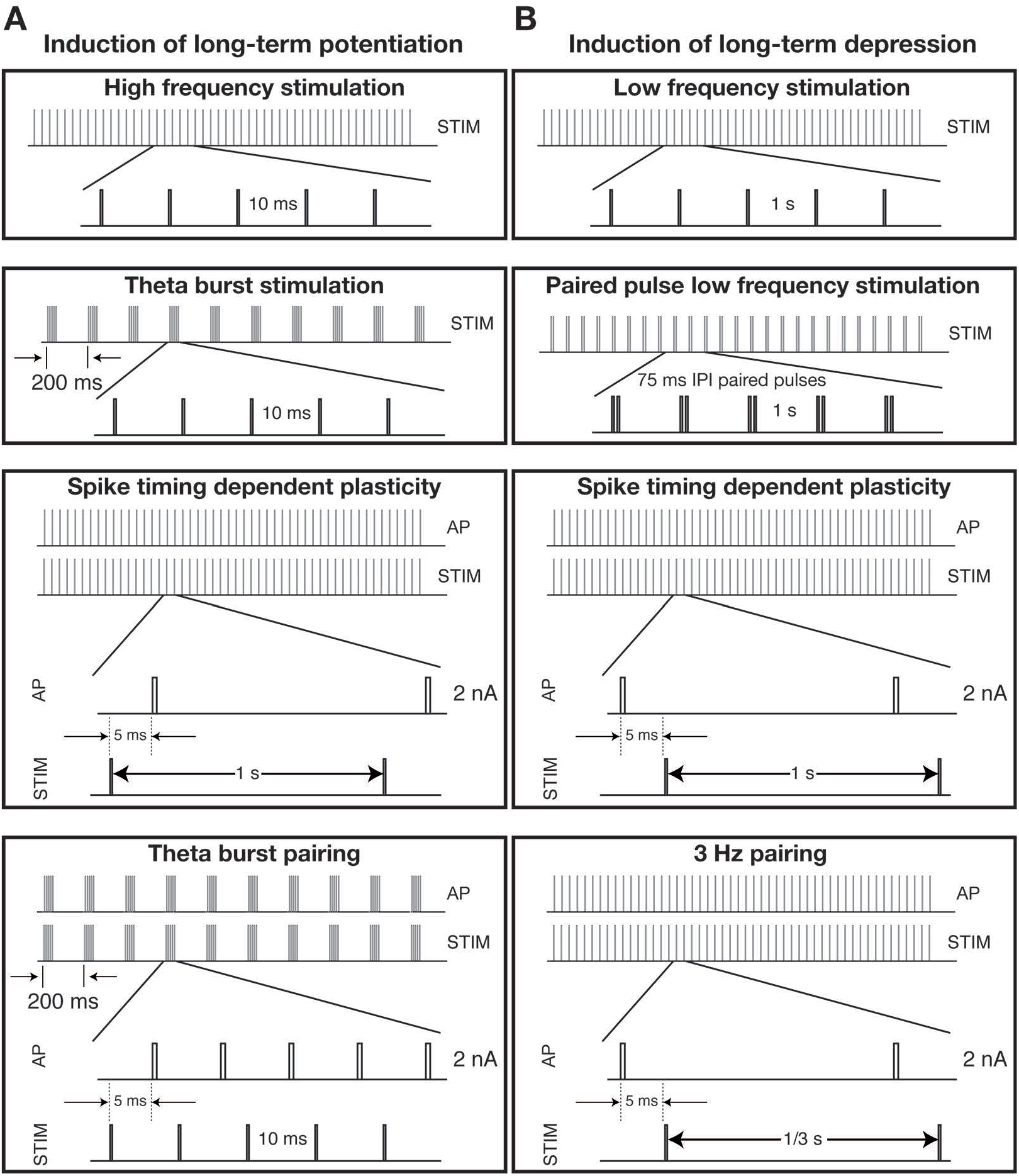
Disparate activity-dependent protocols have been employed for the induction of long-term potentiation or depression in hippocampal synapses. (A–B) Disparate activity-dependent induction protocols yield long-term potentiation (A) or depression (B) in Schaffer collateral synapses connecting CA3 pyramidal neurons to CA1 pyramidal neurons. Individual panels depict cartoon illustrations of induction protocols employed in previous studies (Bi and Poo, 1998; Christie *et al*., 1996; Dudek and Bear, 1992; Huber *et al*., 2000; Larson *et al*., 1986; Magee and Johnston, 1997). AP: action potential; STIM: stimulation leading to postsynaptic potentials; IPI: inter pulse interval. A subset of similar or additional protocols that have been employed in the induction of potentiation or depression in hippocampal synapses may be found here: (Basu *et al*., 2016; Bittner *et al*., 2015; Bittner *et al*., 2017; Bliss and Collingridge, 1993; Bliss and Gardner-Medwin, 1973; Bliss and Lomo, 1973; Chevaleyre *et al*., 2006; Christie *et al*., 1994; Dan and Poo, 2006; Dudek and Bear, 1992, 1993; Dudman *et al*., 2007; Larkman and Jack, 1995; Lynch *et al*., 1983; Lynch *et al*., 1977; Malenka *et al*., 1992; Mulkey and Malenka, 1992; Raymond, 2007; Regehr *et al*., 2009; Staubli and Lynch, 1990; Takahashi and Magee, 2009).

The ability of multiple activity protocols (Fig. 4) to elicit similar levels of synaptic plasticity might be an example of multiple realizability, but it could be argued that this does not constitute an instance of degeneracy, which requires that disparate *structural* components elicit similar function. To address this argument, we refer to established answers for one of the fundamental questions on synaptic plasticity: What is the mechanistic basis for these induction protocols to elicit synaptic plasticity? The influx of calcium into the cytosol is considered as the first step that results in the induction of LTP or LTD (Lynch *et al*., 1983; Malenka *et al*., 1992; Mulkey and Malenka, 1992). Quantitatively, there have been suggestions for the amplitude, spread and kinetics of cytosolic calcium elevation to be specific attributes that translate to the strength and direction of plasticity (Larkman and Jack, 1995; Lisman, 1989; Lisman, 2001; Shouval *et al*., 2002). From this perspective, it may be argued that disparate protocols for inducing LTP (or LTD) result in similar amplitude, spread and kinetics of calcium elevation, thereby resulting in similar strength of LTP (or LTD). With calcium elevation established as a mechanistic basis for the induction of synaptic plasticity, the question of degeneracy here should now focus on the structural basis for eliciting similar elevation in cytosolic calcium.

The mechanisms that govern the strength, spread and kinetics of neuronal calcium are well studied (Augustine *et al*., 2003; Berridge, 1998, 2002, 2006; Berridge *et al*., 2000; Frick *et al*., 2003; Higley and Sabatini, 2012; Jaffe *et al*., 1992; Miyakawa *et al*., 1992; Rizzuto and Pozzan, 2006; Ross, 2012; Sabatini *et al*., 2002; Yasuda *et al*., 2004). Briefly, synergistic interactions between three prominent sets of mechanisms (Fig. 5) regulate cytosolic calcium levels, especially from the perspective of induction of synaptic plasticity. First, the disparate structural components through which calcium ions flow into the cytosol either from the extracellular matrix or from the endoplasmic reticulum (ER). These are typically receptors or channels expressed on the plasma membrane or the ER membrane. The second set is built of disparate mechanisms that alter postsynaptic excitability, which mediates the conversion from synaptic current to synaptic voltage responses. Changes in excitability modulate voltage-levels, which in turn alter calcium influx through voltage-sensitive synaptic receptors or voltage-gated calcium channels. Finally, the expression of calcium-handling mechanisms such as pumps, exchangers and buffers limit the spatiotemporal spread of calcium thereby maintaining specificity of signaling, apart from regulating the strength and kinetics of calcium influx. Thus there are disparate mechanisms that regulate calcium influx, and non-unique combinations of these mechanisms could yield similar strength and kinetics of calcium influx in response to different induction protocols.

**Figure 5.**
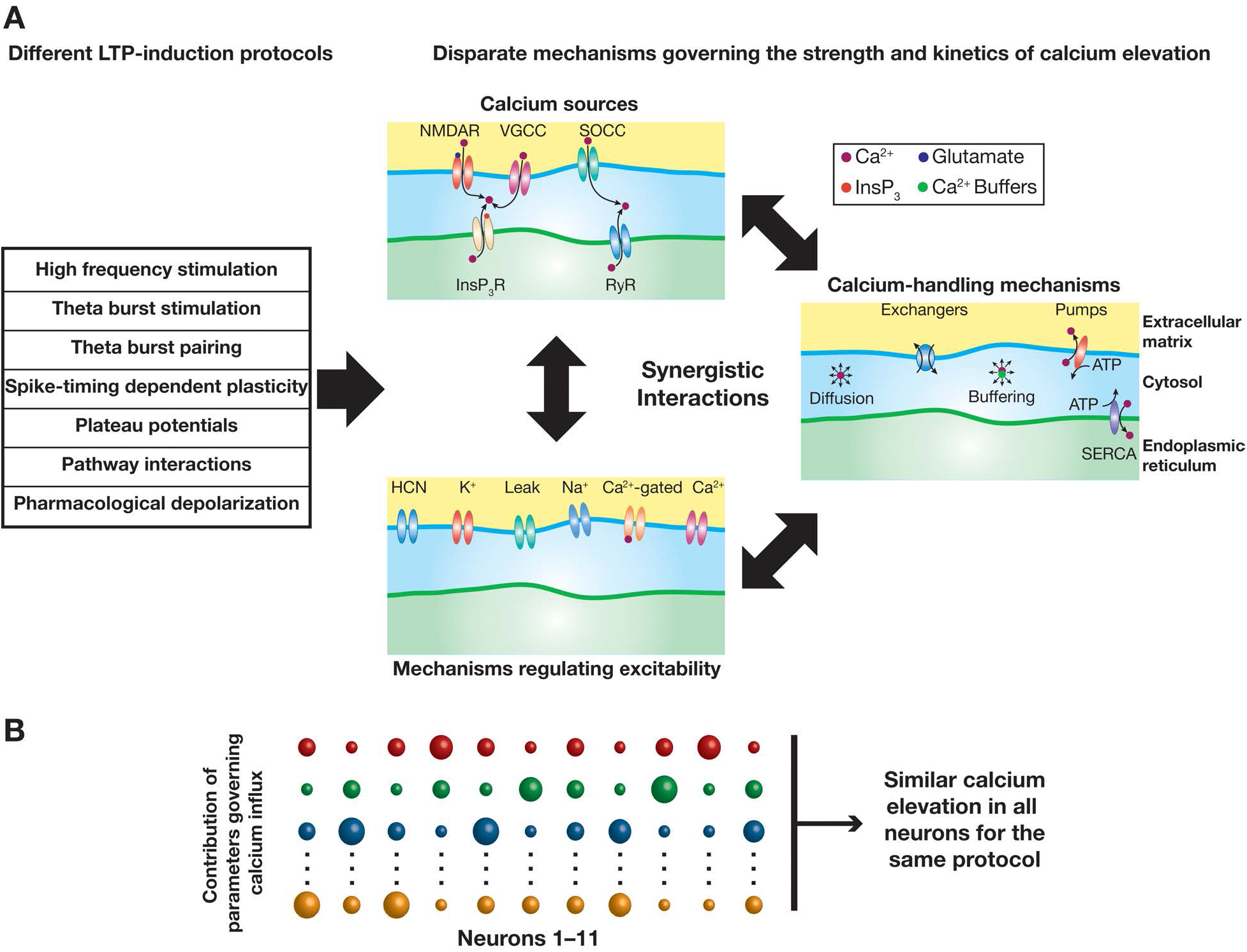
Disparate cellular and molecular mechanisms govern the strength and kinetics of cytosolic calcium influx. (A) Different protocols have been employed for the induction of LTP in hippocampal synapses. Whereas references for the first four of these protocols are provided in Fig. 4, the last three are derived from protocols in these references (Basu *et al*., 2016; Bittner *et al*., 2015; Bittner *et al*., 2017; Dudman *et al*., 2007; Hanse and Gustafsson, 1994; Huang and Malenka, 1993; Huber *et al*., 1995; Lin *et al*., 2008; Otmakhov *et al*., 2004; Roth-Alpermann *et al*., 2006; Takahashi and Magee, 2009). (B) Protocols shown in (A) typically elicit postsynaptic calcium influx through synergistic interactions between disparate constitutive components. Although only postsynaptic components are depicted here, it should be noted that presynaptic components, including excitability-, calcium- and release-regulating mechanisms, also would control the postsynaptic calcium influx through regulation of release dynamics and short-term plasticity. Additionally induction could also be presynaptic. (C) In different neurons, the contribution of different components to achieve similar strength and kinetics of cytosolic calcium influx could be variable. The size of each sphere scales with the quantum of contribution of a given component to cytosolic calcium influx in a given neuron (11 neurons are depicted). Cartoon representations depicted here are drawn from conclusions arrived in previous studies (Anirudhan and Narayanan, 2015; Mukunda and Narayanan, 2017).

Importantly, electrophysiological recordings coupled with pharmacological treatments provide strong lines of evidence that induction of synaptic plasticity could indeed be mediated and regulated by these distinct components. Specifically, there are strong lines of evidence that the induction of bidirectional synaptic plasticity in the hippocampus is mediated by different calcium sources, with certain protocols requiring synergistic activation of multiple calcium sources (Brager and Johnston, 2007; Christie *et al*., 1996; Golding *et al*., 2002; Huber *et al*., 1995; Nishiyama *et al*., 2000; Raymond, 2007). These studies show that plasticity induction is dependent on influx of calcium through NMDA receptors (Christie *et al*., 1996; Collingridge and Bliss, 1987; Collingridge *et al*., 1983; Morris *et al*., 1986; Mulkey and Malenka, 1992; Nishiyama *et al*., 2000; Tsien *et al*., 1996; Wang *et al*., 2003), voltage-gated calcium channels (Brager and Johnston, 2007; Christie *et al*., 1996; Christie *et al*., 1997; Johnston *et al*., 1992; Moosmang *et al*., 2005; Nicholson and Kullmann, 2017; Wang *et al*., 2003), store-operated calcium channels (Baba *et al*., 2003; Garcia-Alvarez *et al*., 2015; Majewski and Kuznicki, 2015; Majewski *et al*., 2016; Prakriya and Lewis, 2015) and receptors on the ER activated by metabotropic receptors on the plasma membrane (Huber *et al*., 2000; Nishiyama *et al*., 2000; Verkhratsky, 2002). Additionally, voltage-gated channels and their auxiliary subunits (Anirudhan and Narayanan, 2015; Brager *et al*., 2013; Chen *et al*., 2006; Chung *et al*., 2009a; Chung *et al*., 2009b; Johnston *et al*., 2003; Jung *et al*., 2008; Kim *et al*., 2007; Lin *et al*., 2008; Lujan *et al*., 2009; Malik and Johnston, 2017; Nolan *et al*., 2004; Sehgal *et al*., 2013; Shah *et al*., 2010; Watanabe *et al*., 2002) have also been shown to critically regulate the strength and direction of synaptic plasticity. Thus, several structural components that mediate or modulate calcium influx into the cytosol have been demonstrated as critical regulators of the induction of synaptic plasticity, both from the qualitative perspective of expression of plasticity and the quantitative perspective of the specific levels of plasticity attained with an induction protocol. Finally, computational modeling has demonstrated that similar synaptic plasticity profiles could be achieved through disparate combinations of channels and receptors (Anirudhan and Narayanan, 2015; Ashhad and Narayanan, 2013; Narayanan and Johnston, 2010; Shouval *et al*., 2002) and is critically dependent on the state of the synapse (Migliore *et al*., 2015). In conjunction with the experimental studies reviewed above, these provide very strong lines of evidence for degeneracy in the induction of synaptic plasticity, where similar levels of calcium influx and analogous synaptic plasticity could be achieved through disparate combinations of parameters that synergistically regulate calcium influx (Fig. 4B).

### 3.4. Degeneracy in signaling cascades that regulate synaptic plasticity

What follows calcium elevation in the process of inducing synaptic plasticity? Once specific strengths and kinetics of calcium influx are achieved as a consequence of induction protocols activating the several disparate mechanisms, is the route to the expression of synaptic plasticity unique? Could multiple mechanisms be activated in response to similar elevations of cytosolic calcium towards achieving specific levels of synaptic plasticity? In other words, is there degeneracy in terms of distinct pathways involving different constitutive components that could link the induction of synaptic plasticity to its expression?

The large body of literature on the signaling cascades involved in synaptic plasticity has presented several lines of evidence that there are several signaling routes, contributing synergistically or differentially, to achieving the translation from the induction of synaptic plasticity to its expression (Fig. 6). Specifically, there is evidence that there are several biochemical species that control synaptic efficacy through a complex network of spatiotemporally interacting signaling cascades (Bhalla, 2014; Bhalla and Iyengar, 1999; Derkach *et al*., 2007; Kennedy, 2000; Kennedy *et al*., 2005; Kholodenko, 2006; Kotaleski and Blackwell, 2010; Larkman and Jack, 1995; Manninen *et al*., 2010; Neves and Iyengar, 2009; Neves *et al*., 2008b; Regehr *et al*., 2009; Weng *et al*., 1999). It is also clear that the dominance of any specific cascade that determines the strength and direction of plasticity is dependent on synaptic state (Migliore *et al*., 2015), the protocol employed (Kandel *et al*., 2014; Mayford *et al*., 2012) and on the spatiotemporal dynamics of changes in the postsynaptic calcium concentration (Berridge, 1998; Korte and Schmitz, 2016; Lisman, 1989; Lisman, 2001; Parekh, 2008; Rizzuto and Pozzan, 2006).

**Figure 6.**
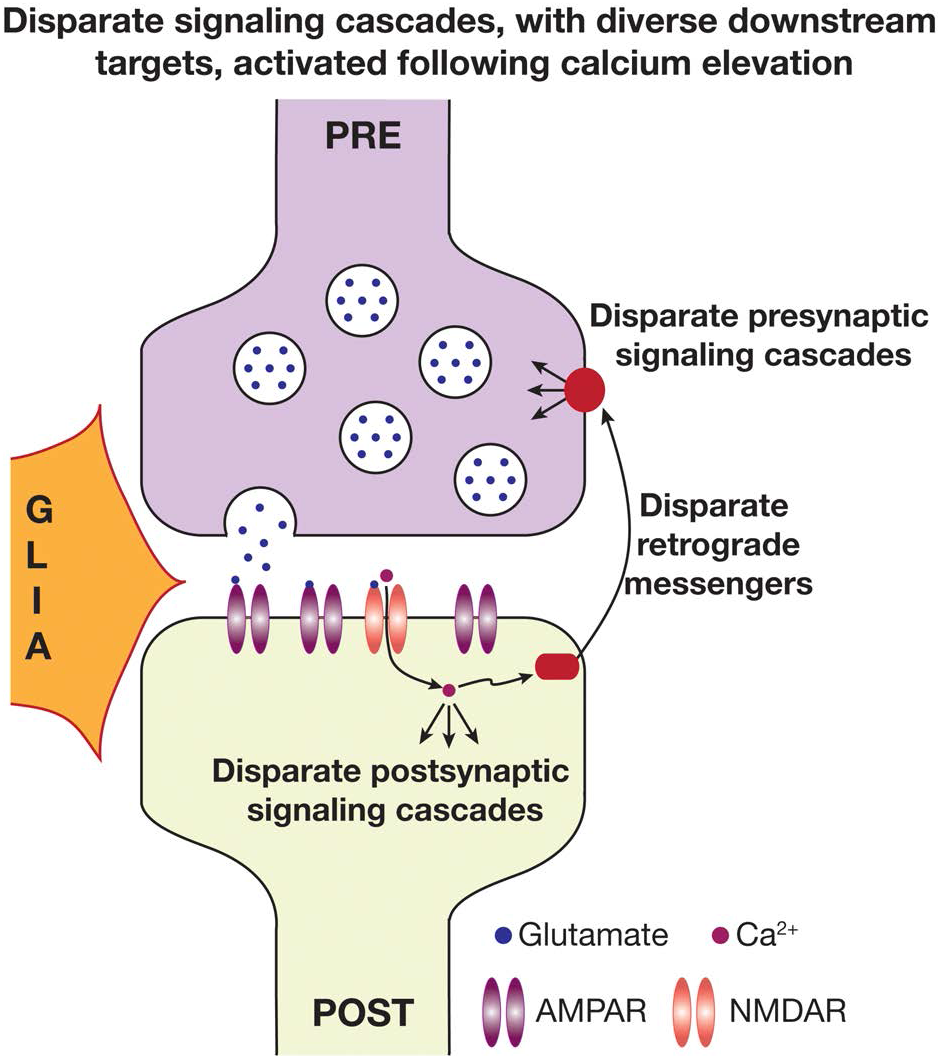
Disparate signaling cascades with diverse downstream targets are activated following postsynaptic calcium elevation. Depicted is a tripartite synapse that includes a presynaptic terminal, a postsynaptic structure and a glial cell. Following the influx of calcium through disparate sources (see Fig. 5; shown here is only NMDAR for simplicity), several pre- and post-synaptic signaling cascades could be activated with very different downstream targets. Retrograde messengers are responsible for intimating the presynaptic terminal about postsynaptic calcium elevation. Illustration incorporates conclusions from previous studies (Bhalla, 2014; Bhalla and Iyengar, 1999; Kotaleski and Blackwell, 2010; Manninen *et al*., 2010; Regehr, 2012; Regehr *et al*., 2009).

The biochemical signaling diversity involved in synaptic plasticity spans both the pre- and post-synaptic sides. The signaling cascades involved in the translation of induction to expression include several enzymes that mediate posttranslational modification of disparate protein substrates, protein synthesis regulators, retrograde messengers, protein trafficking regulators and mechanisms mediating structural plasticity. As a specific example, with reference to the diversity of enzymes that are involved in post-translational modifications resulting in the expression of synaptic plasticity, it has been shown that different protocols for inducing LTP in the Schaffer collateral synapses projecting to CA1 are differentially dependent on different kinases (Kandel, 2001; Kandel *et al*., 2014; Manninen *et al*., 2010; Mayford *et al*., 2012; Raymond, 2007; Soderling and Derkach, 2000). Example kinases are the calcium-calmodulin kinase II, CaMKII (Lisman *et al*., 2002; Lisman *et al*., 2012; Malinow *et al*., 1989; Ouyang *et al*., 1997; Ouyang *et al*., 1999), protein kinase A, PKA (Frey *et al*., 1993; Lin *et al*., 2008; Otmakhova *et al*., 2000; Rosenkranz *et al*., 2009; Woo *et al*., 2003) and mitogen associated protein kinase, MAPK (English and Sweatt, 1997; Rosenkranz *et al*., 2009), which could be activated with the same or different LTP protocols. For instance, the theta-burst pairing protocol activates all of CaMKII, MAPK and PKA (Fan *et al*., 2005; Lin *et al*., 2008; Rosenkranz *et al*., 2009), with very different target substrates involving different channels and receptors (see Sec. 3.6). Additionally the expression of synaptic plasticity, or the substrate for altered synaptic efficacy, could be dependent on several factors (Sec. 3.5), each of which could undergo distinct plasticity with reference to the same activity protocols (Sec. 3.6). Together, the possible combinations of mechanisms that could mediate the translation of plasticity induction protocol to plasticity expression, even for a single synaptic subtype, are numerous. There are also lines of evidence that similar strength and direction of synaptic plasticity could be achieved through the activation of disparate combinations of these mechanisms, providing evidence for the manifestation of degeneracy in the signaling cascades that mediate the transition from plasticity induction to expression.

### 3.5. Degeneracy in the expression of synaptic plasticity

The above analyses establish that hippocampal neurons exhibit degeneracy with reference to the induction of synaptic plasticity and in terms of the mechanisms that mediate the transition from induction to expression. Do these mechanisms act in concert to alter a single target to effectuate the expression of synaptic plasticity? Or are there multiple targets that could be altered to achieve similar strength and direction of synaptic plasticity in a specific synapse?

From the very first study that demonstrated LTP, it has been clear that the protocols employed for inducing synaptic plasticity can recruit different structural components (Bliss and Lomo, 1973):

> “The results suggest that two independent mechanisms are responsible for long-lasting potentiation: (a) an increase in the efficiency of synaptic transmission at the perforant path synapses; (b) an increase in the excitability of the granule cell population.”

Several studies that followed up on this landmark study have now clearly shown that there are disparate routes to achieving synaptic plasticity, even with very similar strength and the same direction of plasticity (Fig. 7). It is now well established that the expression of synaptic plasticity could recruit mechanisms spanning pre- and post-synaptic components, including channels/receptors, morphological features and cytoplasmic constituents on either side (Fig. 7). In other words, different combinations of changes in presynaptic channels/receptors, release mechanisms and postsynaptic channels/receptors could mediate the expression of synaptic plasticity.

**Figure 7.**
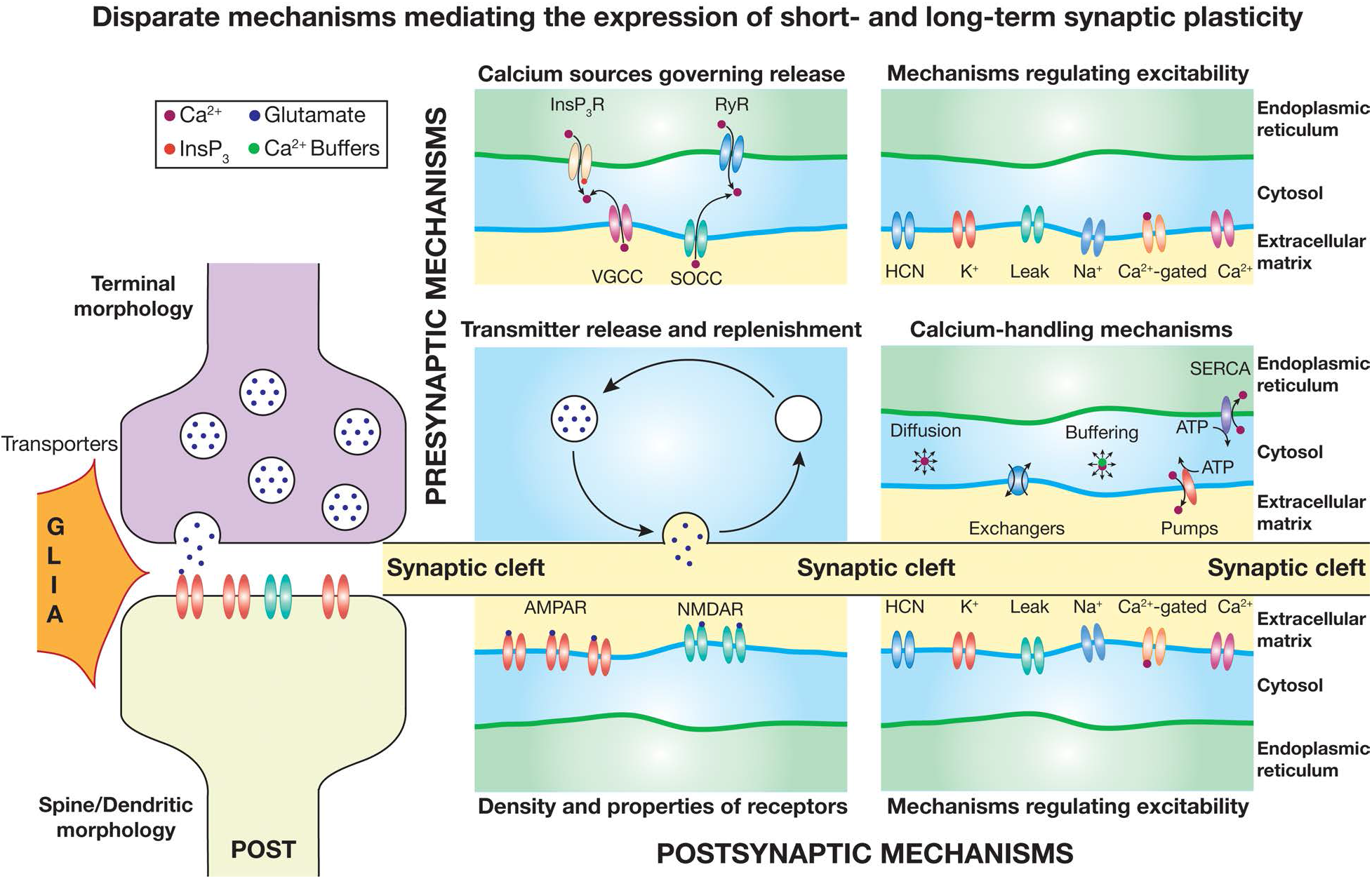
Disparate mechanisms mediate the expression of short- and long-term synaptic plasticity. *Left*, Depicted is a tripartite synapse that includes a presynaptic terminal, a postsynaptic structure and a glial cell. *Right*, Several pre- and post-synaptic mechanisms regulate synaptic strength, and independent or concomitant long-term changes in any of these components would result in the expression synaptic plasticity. Plasticity is known to potentially span all these components and more (Kim and Linden, 2007).

The framework of degeneracy provides an ideal way to reconcile the thorny debates regarding pre- and post-synaptic mechanisms that could mediate synaptic plasticity. Specifically, within this framework, pre- and post-synaptic components would be considered simply as *a subset* (see Sec. 3.6) of the broad repertoire of mechanisms that are available to the neural system to alter towards achieving a specific level of synaptic plasticity or accomplishing an encoding task. Disparate combinations of these components could synergistically contribute to the expression of specific levels of plasticity, at times even with temporal differences in the expression of plasticity in different components. The specific combination of changes that are recruited to mediate plasticity for a chosen protocol or for a given behavioral task would then be state-dependent, critically reliant on the specific calcium sources (Sec. 3.3) and signaling cascades (Sec. 3.4) that were recruited in response to the induction protocol or a behavioral task. In addition to these neuronal components, glial cells, through several mechanisms including gliotransmission or transmitter reuptake and recycling mechanisms, have also been shown to play a critical role in synaptic plasticity (Araque *et al*., 2014; Ashhad and Narayanan, 2016; Halassa *et al*., 2007; Haydon and Carmignoto, 2006; Henneberger *et al*., 2010; Pannasch and Rouach, 2013; Perea and Araque, 2007; Perea *et al*., 2016; Zorec *et al*., 2012), thereby adding another layer of parameters and another set of interactional complexity to the mechanistic basis for synaptic plasticity.

This combinatorial complexity of parameters and associated interactions provide a strong foundation for degeneracy in the emergence of not just the induction and expression of long-term plasticity, but also in the emergence of short-term synaptic plasticity. Specifically, several of the components involved in the induction and expression of long-term plasticity have also been shown to play critical roles in short-term forms of plasticity such as paired pulse facilitation, and on the synaptic filters that they mediate (Atwood *et al*., 2014; Bouchard *et al*., 2003; De Pitta *et al*., 2011; Dittman *et al*., 2000; Emptage *et al*., 2001; Fioravante and Regehr, 2011; Fortune and Rose, 2001; Regehr, 2012; Siegelbaum, 2000; Zucker, 1989, 1999; Zucker and Regehr, 2002). These observations, in conjunction with quantitative computational models have led to the suggestion for the manifestation of degeneracy in the emergence of short-term plasticity profiles and associated synaptic filters (Mukunda and Narayanan, 2017). Specifically, it has been demonstrated that analogous synaptic filters emerge from disparate combinations of presynaptic parameters (Mukunda and Narayanan, 2017). Together, these observations provide clear lines of evidence for the manifestation of degeneracy in short- and long-term forms of synaptic plasticity in the hippocampus.

### 3.6 Degeneracy in the induction and expression of non-synaptic plasticity

It is now widely acknowledged that plasticity protocols and learning paradigms that were once assumed to exclusively recruit or induce synaptic plasticity also induce plasticity in other components (Fig. 8), in a manner that could either be localized or global. Similar to the study of synaptic plasticity, specific activity protocols (most of which are similar, if not identical, to synaptic plasticity protocols) are employed to assess plasticity in other protein molecules and structural changes. Plasticity in voltage-gated ion channels and other neuronal components that result in changes to neuronal intrinsic properties have been dubbed as *intrinsic plasticity*, and is known to occur in the hippocampus with reference to most activity-dependent protocols employed for inducing synaptic plasticity (Brager and Johnston, 2007; Chung *et al*., 2009a; Chung *et al*., 2009b; Fan *et al*., 2005; Frick and Johnston, 2005; Frick *et al*., 2004; Johnston *et al*., 2003; Johnston and Narayanan, 2008; Kim and Linden, 2007; Lin *et al*., 2008; Losonczy *et al*., 2008; Magee, 2000; Mozzachiodi and Byrne, 2010; Narayanan and Johnston, 2007, 2008, 2012; Nelson and Turrigiano, 2008; Remy *et al*., 2010; Sjostrom *et al*., 2008; Spruston, 2008; Wang *et al*., 2003; Zhang and Linden, 2003). Although it is generally assumed that intrinsic plasticity refers only to global changes in intrinsic *excitability*, it is important to recognize that intrinsic plasticity encompasses *all* intrinsic properties that are mediated by neuronal constitutive components (Llinas, 1988; Marder, 2011; Marder *et al*., 1996; Marder and Goaillard, 2006), including neuronal spectral selectivity conferred by specific sets of ion channels (Das *et al*., 2017; Hutcheon and Yarom, 2000) and calcium wave propagation mediated by receptors on the endoplasmic reticulum (Ross, 2012). These distinct intrinsic properties, including excitability, have been shown to undergo bidirectional changes in a manner that is local to specific neuronal locations or is global spanning all locations (Brager and Johnston, 2007; Das *et al*., 2017; Johnston and Narayanan, 2008; Narayanan *et al*., 2010; Narayanan and Johnston, 2007, 2008).

**Figure 8.**
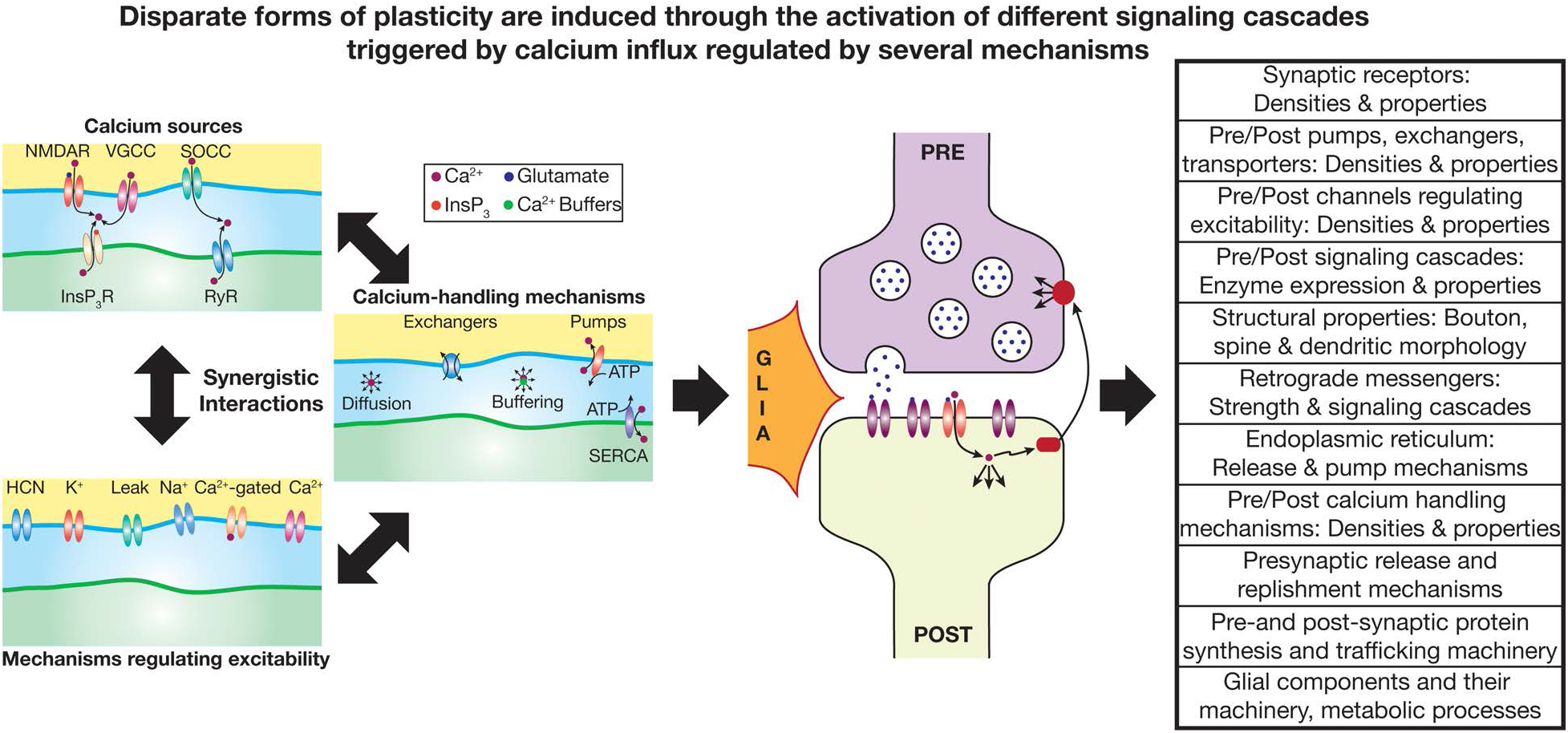
Disparate forms of synaptic and non-synaptic plasticity are induced through the activation of different signaling cascades triggered by calcium influx regulated by several mechanisms, resulting in multiscale degeneracy in plasticity induction through expression. *Left*, Synergistic interactions between several components results in cytosolic calcium influx following plasticity induction through activity protocols or behavioral experience of pathological insults. *Center*, Disparate signaling cascades with diverse downstream targets are activated following postsynaptic calcium elevation. *Right*, The activation of signaling cascades and their impact on their targets are not just limited to synaptic components, but span a large span of neuronal and network components. Several forms of synaptic and non-synaptic plasticity express concomitantly in response to the same protocols or perturbations (Beck and Yaari, 2008; Johnston *et al*., 2016; Kim and Linden, 2007; Narayanan and Johnston, 2012).

As the protocols employed for inducing non-synaptic (including intrinsic and structural) plasticity are at most instances identical to synaptic plasticity induction protocols, the broad mechanisms involved in the induction and in the translation of induction to expression are very similar to those for synaptic plasticity (Fig. 8). Specifically, induction of intrinsic plasticity requires influx of cytosolic calcium with different kinetics and strengths of calcium translating to distinct strengths and directions of intrinsic plasticity (Brager and Johnston, 2007; Fan *et al*., 2005; Huang *et al*., 2005; Sjostrom *et al*., 2008; Wang *et al*., 2003). The components that mediate calcium entry for synaptic plasticity also mediate calcium entry for non-synaptic plasticity, including NMDA receptors (Chung *et al*., 2009a; Chung *et al*., 2009b; Engert and Bonhoeffer, 1999; Fan *et al*., 2005; Frick *et al*., 2004; Huang *et al*., 2005; Lin *et al*., 2008; Losonczy *et al*., 2008; Matsuzaki *et al*., 2004; Nagerl *et al*., 2004; Narayanan and Johnston, 2007; Tonnesen *et al*., 2014; Wang *et al*., 2003), voltage-gated calcium channels (Chung *et al*., 2009a; Chung *et al*., 2009b; Lin *et al*., 2008; Wang *et al*., 2003) and receptors on the ER (Ashhad *et al*., 2015; Brager and Johnston, 2007; Brager *et al*., 2013; Clemens and Johnston, 2014; Kim *et al*., 2017; Narayanan *et al*., 2010). This implies that the arguments (Secs. 3.3–3.4) placed about synergistic interactions between different calcium sources and about degeneracy in the induction of synaptic plasticity extends to the induction of non-synaptic plasticity as well (Fig. 8).

As a direct consequence of the similarity in the protocols employed in inducing synaptic and intrinsic plasticity, the downstream mechanisms that mediate the translation from induction of non-synaptic plasticity to its expression are also similar (Shah *et al*., 2010) to those that mediate a similar transition in synaptic plasticity (Fig. 8). Several signaling cascades that are present on the pre- and post-synaptic sides mediate this translation, with retrograde messengers acting as mechanisms that signal the elevation of postsynaptic calcium to the presynaptic terminals. Specifically, the same set of enzymes and messengers that mediate synaptic plasticity also mediate non-synaptic plasticity (Fig. 8). Examples to this equivalence include non-synaptic forms of plasticity that are mediated by CaMKII (Fan *et al*., 2005; Huang *et al*., 2005; Lujan *et al*., 2009; Matsuzaki *et al*., 2004; Wang and Wagner, 1999), PKA (Lin *et al*., 2008; Narayanan *et al*., 2010; Rosenkranz *et al*., 2009) and MAPK (Rosenkranz *et al*., 2009; Yuan *et al*., 2002). However, there could be dissociation between the mechanisms that are involved in the translation to the expression of different forms of plasticity that are consequent to the *same* induction protocol, where different enzymes and messengers mediate different forms of plasticity (Brager and Johnston, 2007; Fan *et al*., 2005; Lin *et al*., 2008; Rosenkranz *et al*., 2009; Wang *et al*., 2003). As mentioned earlier (Sec. 3.5), the expression of plasticity in synapses could be mediated by plasticity in voltage-gated calcium channels that are expressed in the presynaptic terminal, mediated by retrograde messengers and presynaptic signaling cascades, or by change in mechanisms that alter postsynaptic excitability, thus blurring the distinction between synaptic and certain forms of non-synaptic plasticity.

Following the activation of different signaling cascades, akin to the expression of synaptic plasticity, several molecular processes, including synthesis, trafficking and post-translational modification of the several membrane and cytosolic proteins, mediate the final step towards the expression of distinct forms of non-synaptic plasticity (Fig. 8). The mechanisms behind the trafficking of several ion channels have been studied (Cusdin *et al*., 2008; Jensen *et al*., 2011; Lai and Jan, 2006; Lau and Zukin, 2007; Lujan *et al*., 2009; Shah *et al*., 2010; Vacher *et al*., 2008; Wenthold *et al*., 2003), and it is now clear that plasticity is ubiquitous (Kim and Linden, 2007). In addition to these changes in cytosolic and membrane proteins, it has been shown that hippocampal spines undergo continuous structural changes, apart from demonstrations of distinct forms of structural plasticity in spines, dendrites and axons (Attardo *et al*., 2015; Chen *et al*., 2014; Emoto, 2011; Engert and Bonhoeffer, 1999; Ghiretti and Paradis, 2014; Grubb and Burrone, 2010a, b; Grubb *et al*., 2011; Ikegaya *et al*., 2001; Johnston *et al*., 2016; Luo and O’Leary, 2005; Matsuzaki *et al*., 2004; Nagerl *et al*., 2004; Tonnesen *et al*., 2014; Yuste and Bonhoeffer, 2001). Finally, the dynamics associated with the various glial functions and their interactions with neuronal and metabolic pathways could also undergo changes in response to behavioral experiences and activity (Araque *et al*., 2014; Baumann and Pham-Dinh, 2001; Fields, 2010; Halassa and Haydon, 2010; Haydon and Carmignoto, 2006; Khakh and Sofroniew, 2015; Pannasch and Rouach, 2013; Perea *et al*., 2016; Sierra *et al*., 2014; Volterra *et al*., 2014). It is therefore clear that there is no escape from the conclusion that activity-or experience-or pathology-dependent plasticity does not confine itself to a few constitutive components, but is rather expansive and even ubiquitous (Kim and Linden, 2007). There are considerable overlaps in the mechanisms that mediate the induction and expression of these forms of plasticity, and many-to-one and one-to-many mappings between the induction protocol (or behavioral experience) and achieving specific levels of plasticity in specific components (Fig. 8).

In summary, the lines of evidence provided above point to ample evidence for degeneracy in the process of their induction and expression of different forms of plasticity and their combinations, both in terms of their individual strengths and directions. This also implies that the same functional changes could be achieved through distinct combinations of plasticity mechanisms, thus pointing to a further dissociation between functional homeostasis and the plasticity mechanisms that yielded it. In other words, functional equivalence in terms of transition from one state to another does not necessarily translate to plasticity equivalence (where the route taken to achieve the transition is always identical). An important class of plasticity models has recognized the ubiquitous nature of plasticity, with models built within this framework of plasticity degeneracy. These models account for concomitant changes in multiple components, also accounting for disparate combinations of plasticity resulting in similar functional outcomes, rather than assuming plasticity equivalence in the face of functional equivalence (Abbott and LeMasson, 1993; Anirudhan and Narayanan, 2015; LeMasson *et al*., 1993; Mukunda and Narayanan, 2017; O’Leary *et al*., 2013; O’Leary *et al*., 2014; Siegel *et al*., 1994; Srikanth and Narayanan, 2015). Future theoretical and experimental investigations into hippocampal plasticity should therefore account for the truly ubiquitous nature of plasticity in designing their experiments and addressing outstanding questions, rather than assuming that plasticity is confined to one single component or the other (Bhalla, 2014; Kim and Linden, 2007).

### 3.7 Degeneracy in metaplasticity and in maintaining stability of learning

Hebbian synaptic plasticity is inherently unstable. In the absence of concomitant homeostatic mechanisms, Hebbian plasticity would result in runaway excitation (Fig. 9). Several theories and mechanisms have been proposed as a means to avoid this runaway excitation (Abbott, 2003; Abraham and Robins, 2005; Korte and Schmitz, 2016; Miller and MacKay, 1994; Nelson and Turrigiano, 2008; Turrigiano, 2007, 2011; Turrigiano, 1999, 2008, 2017; Turrigiano and Nelson, 2000; van Rossum *et al*., 2000; Zenke *et al*., 2017). A prominent theme that spans several such stability theories is metaplasticity (Abraham, 2008; Abraham and Bear, 1996; Abraham and Tate, 1997; Hulme *et al*., 2013), where the profile of plasticity concomitantly changes with the induction of plasticity (Fig. 9A–B). An extremely useful mathematical treatise that has helped in the understanding metaplasticity and stability, especially for synaptic plasticity profiles in the hippocampus, is the Bienenstock-Cooper-Munro (BCM) rule (Bienenstock *et al*., 1982; Cooper and Bear, 2012; Shouval *et al*., 2002; Yeung *et al*., 2004). This is despite the observation that the BCM framework and the synaptic plasticity framework in hippocampal synapses are not completely analogous to each other (Cooper *et al*., 2004). It should also be noted that not all synapses follow a BCM-like synaptic plasticity profile, and therefore a stability theory dependent on this rule is not generalizable to all synapses (Abbott and Nelson, 2000; Jorntell and Hansel, 2006).

**Figure 9.**
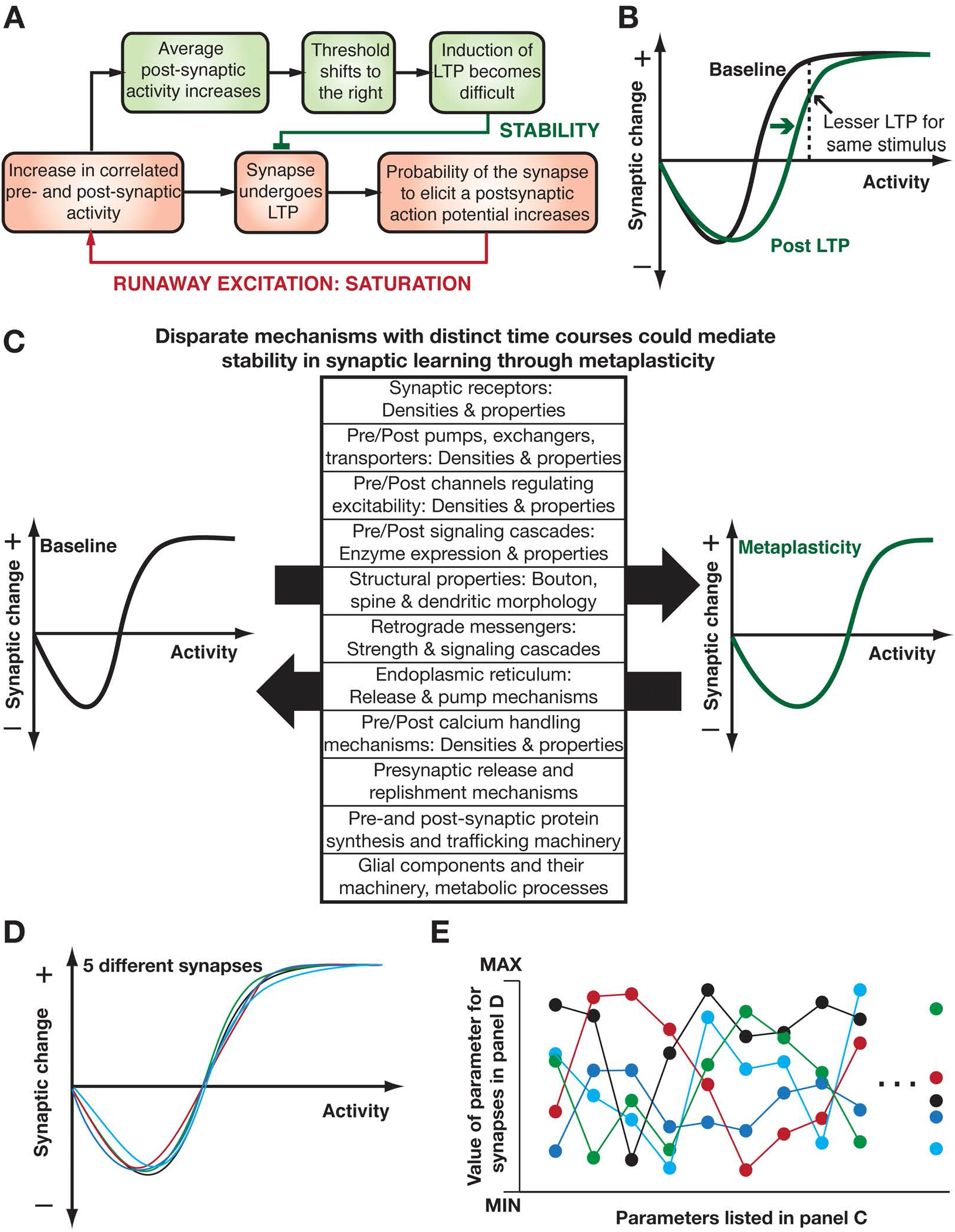
Disparate mechanisms with distinct time courses could mediate stability in synaptic learning through metaplasticity. (A–B) Hebbian synaptic plasticity is inherently unstable leading to runaway excitation in synaptic structure (A; orange boxes). The Bienenstock-Cooper–Munro (BCM) rule envisages the existence of a sliding threshold mechanism (B) which provides a negative feedback loop (B; green boxes) that would preclude runaway excitation by altering the rules for plasticity. Alteration of plasticity rules has been referred to as metaplasticity in the literature (Abraham and Bear, 1996; Bienenstock *et al*., 1982; Cooper and Bear, 2012). (C) Bidirectional metaplasticity could be mediated by any of the several mechanisms discussed in Fig. 7–8 with reference to the expression of synaptic and non-synaptic plasticity. (D–E) Similar plasticity profiles (D) could be achieved through disparate combinations of constituent parameter values (E). Cartoon illustrations are derived from conclusions drawn in previous studies (Abraham, 2008; Abraham and Bear, 1996; Anirudhan and Narayanan, 2015; Hulme *et al*., 2013; Sehgal *et al*., 2013).

Although the utility of BCM-like synaptic rule in understanding stability in synaptic learning has been invaluable, the exact mechanisms that mediate the sliding modification threshold and the consequent metaplasticity has remained an open question. Several mechanisms (Fig. 9C) involving changes in morphological characteristics, several receptors, ion channels and signaling cascades have been proposed as candidates for this role (Abraham, 2008; Abraham and Bear, 1996; Abraham *et al*., 2001; Abraham and Tate, 1997; Anirudhan and Narayanan, 2015; Bear, 2003; Bear *et al*., 1987; Cooper and Bear, 2012; Hulme *et al*., 2013; Kalantzis and Shouval, 2009; Narayanan and Johnston, 2010; Philpot *et al*., 2003; Philpot *et al*., 2001; Sehgal *et al*., 2013; Triesch, 2007). As any change in mechanisms that regulate the induction or expression of synaptic plasticity would result in a change in plasticity profiles (Sec. 3.3–3.5), it is not surprising that mechanisms that regulate synaptic plasticity are candidate mechanisms that mediate metaplasticity. Similar to the argument placed with reference to the mechanisms that mediate the expression of synaptic plasticity, the framework of degeneracy provides an elegant solution to the question on which of these is *the* mechanism that mediates the sliding modification threshold within a BCM-like plasticity framework. It offers reconciliation to this conundrum by suggesting that disparate combinations of these distinct mechanisms could result in similar plasticity profiles (Fig. 9D–E), thereby suggesting degeneracy in the emergence of metaplasticity and stability in synaptic learning (Anirudhan and Narayanan, 2015). Finally, it was traditionally assumed that stability and homeostatic mechanisms are slower compared to the encoding mechanisms. However, there are several lines of theoretical and experimental evidence, spanning several synaptic and intrinsic components as candidate mechanisms, for *concurrent* emergence of encoding, stability *and* activity homeostasis. These lines of evidence also argue for prominent advantages when encoding, homeostasis and stability mechanisms are *concurrent* (Anirudhan and Narayanan, 2015; Honnuraiah and Narayanan, 2013; Ibata *et al*., 2008; Jedlicka *et al*., 2015; Johnston and Narayanan, 2008; Narayanan and Johnston, 2007, 2010; Nelson and Turrigiano, 2008; O’Leary *et al*., 2014; Triesch, 2007; Turrigiano, 2011; Turrigiano, 2008, 2017; Zenke *et al*., 2017).

Within the framework of degeneracy, the goal of *concomitantly* achieving encoding-driven plasticity, activity homeostasis and stable learning is achieved through disparate combinations of synaptic, intrinsic, glial and structural plasticity. With abundant experimental evidence for plasticity in each of these different components occurring in an activity-or experience-dependent manner (Sec. 3.6), it is important that the analyses of stable learning broaden their focus beyond the narrow realm of stable *synaptic* learning. The current theories implicitly or explicitly assume that encoding is driven by synaptic plasticity, with several mechanisms contributing to the stability of this synaptic learning system. The metaplasticity framework also largely focuses on plasticity of *synaptic* plasticity profiles, although the mechanisms that mediate several forms of plasticity overlap with each other (Sec. 3.6). Future frameworks should therefore analyze concomitant learning *and* stability as a consequence of disparate forms of plasticity, also assessing *metaplasticity of intrinsic, glial and structural plasticity*. While plasticity in synaptic structures form *a component* of learning and stability, given the abundant lines of experimental evidence on ubiquitous plasticity, it is extremely critical that learning and stability theories broaden their horizon to encompass all forms of plasticity and degeneracy therein.

### 3.8. Degeneracy in the generation and regulation of local field potentials

Extracellular field recordings are useful readouts of network activity in a given brain region. Local field potentials (LFP), the low pass filtered version of field recordings have traditionally been thought to provide information about afferent synaptic activity. LFPs recorded from within the hippocampus exhibit signature activity patterns that are dependent on the behavioral state of the animal. For instance, they manifest strong oscillations in the theta frequency range (4–10 Hz) during exploratory behavior and during rapid eye moment (REM) sleep, and show characteristic sharp-wave ripple patterns during rest and non-REM sleep. These distinct activity patterns have been postulated to serve specific functions such as in the consolidation of memory and in neural encoding of space (Buzsaki, 1986, 1989, 2002, 2006, 2015; Buzsaki and Moser, 2013; Colgin, 2013; English *et al*., 2014; Grosmark *et al*., 2012; Hartley *et al*., 2014; Lisman and Jensen, 2013; Mizuseki and Buzsaki, 2014; Montgomery *et al*., 2008; Moser *et al*., 2008; Moser *et al*., 2015; Tononi and Cirelli, 2006; Wilson and McNaughton, 1994; Ylinen *et al*., 1995a; Ylinen *et al*., 1995b).

Although these signature patterns of extracellular events manifest as repeating motifs, there are strong lines of theoretical and experimental evidence that they emerge from very disparate structures. For instance, theta oscillations in the hippocampus have shown to be afferent from two reciprocally connected subcortical nuclei that act as pacemakers, the medial septum-diagonal band of Broca and the supramammillary region. Apart from these two subcortical nuclei, inputs from entorhinal cortex and CA3 also play an important role in the generation of theta oscillations in the hippocampus. Furthermore, theoretical modeling and *in vitro* data also suggest that an intact hippocampus could sustain theta oscillations on its own in a manner that is dependent on intra-hippocampal excitatory and inhibitory synaptic connections (Buzsaki, 2002, 2006; Colgin, 2013, 2016; Goutagny *et al*., 2009; Kamondi *et al*., 1998; Traub *et al*., 1989). A similar analysis, in terms of disparate underlying sources and mechanisms, holds for gamma frequency oscillations that are observed in the hippocampus as well (Buzsaki and Wang, 2012; Colgin, 2016; Colgin and Moser, 2010; Csicsvari *et al*., 2003; Wang, 2010; Wang and Buzsaki, 1996). In addition, apart from synaptic contributions to the LFPs, it is now clear that return transmembrane currents from sub- and supra-threshold somatodendritic ion channels also alter the LFP in terms of their frequency content, amplitude and phase (Buzsaki *et al*., 2012; Einevoll *et al*., 2013; Ness *et al*., 2016; Reimann *et al*., 2013; Schomburg *et al*., 2012; Sinha and Narayanan, 2015; Taxidis *et al*., 2015). In addition, several mechanisms such ephaptic coupling, heterogeneous extracellular resistivity, glial and axonal transmembrane mechanisms also contribute and regulate local field potentials, resulting in a complexity that spans almost all parameters of the local network (Anastassiou and Koch, 2015; Buzsaki *et al*., 2012; Einevoll *et al*., 2013; Kajikawa and Schroeder, 2011; Katzner *et al*., 2009; Linden *et al*., 2011).

From the complexity involved in the generation and regulation of hippocampal LFPs, with several brain regions and several constitutive network components contributing to their emergence, it is easy to discern that similar LFP patterns could be achieved through non-unique combinations of disparate components. Irrespective of whether it is the manifestation of an oscillatory pattern in a given frequency range (Buzsaki, 2002; Buzsaki and Wang, 2012; Colgin, 2013; Colgin and Moser, 2010; Csicsvari *et al*., 2003), or the emergence of sharp wave ripples (Buzsaki, 2015; English *et al*., 2014; Taxidis *et al*., 2015), or the emergence of resonance in the LFP power spectral density (Ness *et al*., 2016), or achieving a given phase of single-neuron firing with reference to an LFP oscillation (Sinha and Narayanan, 2015), the routes are several and involve several disparate structural components. Thus, there is evidence for degeneracy in the mechanisms that mediate and regulate local field potentials, implying that extreme caution should be exercised in making one-to-one relationships between constitutive components and specific aspects of LFP recordings (Anastassiou and Koch, 2015; Buzsaki *et al*., 2012; Einevoll *et al*., 2013; Kajikawa and Schroeder, 2011; Katzner *et al*., 2009; Linden *et al*., 2011).

### 3.9. Degeneracy in neural coding

A particularly thorny debate that has spanned decades is about the codes employed by neurons in encoding their inputs. The crux of the debate has been about whether neurons encode information in the rate of or in the precise timing of action potential firing (Buzsaki *et al*., 2013; Engel *et al*., 2001; Engel and Singer, 2001; Fries *et al*., 2007; Gallistel, 2017; Jaramillo and Kempter, 2017; London *et al*., 2010; Panzeri *et al*., 2017; Shadlen and Newsome, 1994, 1995, 1998; Singer *et al*., 1997; Softky, 1994; Softky, 1995). Arguments against temporal coding have raised questions about the ability of neurons to perform millisecond-or-submillisecond coincidence detection that is essential for decoding a temporal code, about the relevance of precise timing in the face of noise and variability in neuronal responses to identical stimuli and about the ability of neuronal networks to reliably propagate synchronous firing (London *et al*., 2010; Panzeri *et al*., 2017; Shadlen and Newsome, 1994, 1998). Counterarguments have relied on the demonstration of millisecond-or-submillisecond coincidence detection in active dendritic structures, on the dependence of synchrony propagation on neuronal intrinsic properties *and* input structure and on the existence of temporally precise cell assemblies that could mitigate the overall background noise in decoding the precise timing of inputs (Buzsaki, 2010; Buzsaki *et al*., 2013; Das and Narayanan, 2015, 2017; Diesmann *et al*., 1999; Engel *et al*., 2001; Engel and Singer, 2001; Fries *et al*., 2007; Golding and Oertel, 2012; Hong *et al*., 2012; Pastalkova *et al*., 2008; Reyes, 2003; Singer *et al*., 1997; Softky, 1994).

The expression of *coding* degeneracy in the cellular and network scales (Leonardo, 2005), in terms of the ability of disparate structural components to elicit similar input-output characteristics, is clear from the lines of evidence presented earlier (Sec. 2.2). In addition, employing electrophysiological recordings and computational models to assess subthreshold resonance and spike triggered average (STA) of model neurons, it has been shown that hippocampal pyramidal neurons are selective to different input features (including spectral features and temporal coincidence of inputs) depending on the dendritic location of their inputs. This location-dependent feature encoding is mediated by ion channel expression profiles, and could be achieved through disparate combinations of different ion channel expression profiles (Das and Narayanan, 2014, 2015, 2017; Das *et al*., 2017; Narayanan and Johnston, 2007, 2012; Rathour *et al*., 2016; Rathour and Narayanan, 2012a, 2014). Given the well-established strong relationship between STA and types of coding (Ratte *et al*., 2013), this location-dependent scenario argues for location-dependent forms of coding. Specifically, the soma and proximal dendrites showing class I STA (integrator) and the distal dendrites manifesting class II STA (coincidence detector) as a consequence of the differential expression of different channels (Das and Narayanan, 2015). Therefore, it seems reasonable to postulate that the proximal and distal regions are respectively geared towards rate and temporal coding, with this location-dependent differential coding strategy extending to cortical and hippocampal neurons (Branco and Hausser, 2010, 2011; Das and Narayanan, 2015). Finally, behaviorally-driven neuromodulatory inputs and activity-dependent plasticity could markedly alter the operating mode and the class of excitability of compartments of a single neuron, and the type of coding employed by a neuron is dependent not just on its operating mode but also the specific characteristics of the input. Thus, even from the perspective of encoding strategies *within* a single neuron, the arguments that pitch rate coding *against* temporal coding are oversimplifying the complexity of neural encoding and decoding. Instead, there are broad lines of evidence pointing to a hybrid rate/temporal coding system that encompasses degeneracy by achieving encoding goals through disparate combinations of several cellular and network components in a manner that is strongly dependent on several spatiotemporal aspects of neuronal and behavioral state (Das and Narayanan, 2014, 2015; Das *et al*., 2017; Diesmann *et al*., 1999; Lee and Dan, 2012; Marder, 2012; Marder and Thirumalai, 2002; Ratte *et al*., 2013).

With reference to neural codes for features of the external environment, the coding of spatial location of animal in the hippocampus is an ideal instance of hybrid encoding schema that expresses degeneracy. Unlike the argument for rate *vs*. temporal coding that seems to drive the narrative otherwise (Buzsaki *et al*., 2013; Engel *et al*., 2001; Engel and Singer, 2001; Fries *et al*., 2007; Gallistel, 2017; Jaramillo and Kempter, 2017; London *et al*., 2010; Panzeri *et al*., 2017; Shadlen and Newsome, 1994, 1995, 1998; Singer *et al*., 1997; Softky, 1994; Softky, 1995; Srivastava *et al*., 2017), hippocampal physiologists have concurred on the existence of dual/hybrid encoding schema for place-specific encoding. Specifically, place cells in the hippocampus elicit higher rates of firing when the animal enters a specific place field. In conjunction, the phase of action potential firing of place cells with reference to the extracellular theta rhythm also advances as a function of spatial location of the animal within the place field. Thus, hippocampal place cells employ a dual code of firing rate *and* phase of firing (temporal coding involving the precise timing of action potential firing) to represent spatial location of the animal (Ahmed and Mehta, 2009; Buzsaki and Moser, 2013; Derdikman and Moser, 2010; Hartley *et al*., 2014; Harvey *et al*., 2009; Huxter *et al*., 2003; Lisman, 2005; Lisman and Jensen, 2013; Mehta *et al*., 2002; Moser *et al*., 2008; Moser *et al*., 2015; O’Keefe, 1976, 1979; O’Keefe and Burgess, 1999, 2005; O’Keefe *et al*., 1998; O’Keefe and Conway, 1978; O’Keefe and Recce, 1993; Skaggs *et al*., 1996). In certain cases, it has been shown that the two coding schema act independent of each other and could act as the fail-safe mechanisms for each other (Aghajan *et al*., 2015; Huxter *et al*., 2003).

Whereas these lines of evidence make a case for employing disparate coding schemas in encoding the same input, the case for disparate mechanisms involved in encoding and maintaining the rate and temporal codes is also strong. Specifically, the role of afferent synaptic drive, local inhibition, several ion channels and receptors, dendritic spikes, spatiotemporal interactions between somatodendritic channels and receptors, and plasticity in each of these components have all been implicated in the emergence and maintenance of these codes (Bittner *et al*., 2015; Danielson *et al*., 2016; Geisler *et al*., 2010; Geisler *et al*., 2007; Grienberger *et al*., 2017; Harvey *et al*., 2009; Lee *et al*., 2012; Losonczy *et al*., 2010; Magee, 2001; Nakashiba *et al*., 2008; Nakazawa *et al*., 2004; Nolan *et al*., 2004; Royer *et al*., 2012; Sheffield and Dombeck, 2015; Skaggs *et al*., 1996; Tsien *et al*., 1996; Wills *et al*., 2005). In addition, there are lines of experimental evidence that suggest that subthreshold afferent synaptic inputs from several place fields arrive onto a single place cell, and that a silent cell could be converted to a place cell for *any* of these place fields by an appropriate plasticity-inducing stimulus (Bittner *et al*., 2015; Lee *et al*., 2012), suggesting that disparate cells could achieve the same function of encoding a given spatial location. The expression profiles of several channels and receptors control the overall excitability of a neuron (Sec. 2.2), and there are several mechanisms that regulate the phase of intracellular voltage oscillations with reference to an external reference or to the overall afferent current (Geisler *et al*., 2010; Geisler *et al*., 2007; Harvey *et al*., 2009; Narayanan and Johnston, 2008; Rathour *et al*., 2016; Rathour and Narayanan, 2012a, 2014; Sinha and Narayanan, 2015; Skaggs *et al*., 1996). Together, these studies point to the possibility that similar rate *and* phase spatial codes in a neuron could be achieved through disparate combinations of constituent components, and several neurons could encode for the same place field with distinct combinations of these mechanisms. Future studies could further explore the manifestation of degeneracy in spatial coding in the hippocampus, focusing on the hybrid code involving rate as well as phase encoding of input features.

### 3.10. Degeneracy in learning and memory

Behavior emerges as a consequence of coordinated activity of multiple brain regions in conjunction with sensory and motor systems (Bennett and Hacker, 2003; Jazayeri and Afraz, 2017; Krakauer *et al*., 2017; Tytell *et al*., 2011; Vetere *et al*., 2017). The hippocampus has been implicated in several forms of spatial and non-spatial learning, with strong links to episodic memory (Anderson *et al*., 2007; Bird and Burgess, 2008; Bliss and Collingridge, 1993; Bunsey and Eichenbaum, 1996; Lynch, 2004; Marr, 1971; Martin *et al*., 2000; Martinez and Derrick, 1996; Mayford *et al*., 2012; Morris, 1989; Morris *et al*., 1986; Morris *et al*., 1982; Moser *et al*., 2015; Nakazawa *et al*., 2004; Neves *et al*., 2008a; Rajasethupathy *et al*., 2015; Scoville and Milner, 1957; Squire *et al*., 2004; Whitlock *et al*., 2006).

The quest for *the* mechanistic basis for learning and memory in the hippocampus has spanned several decades, especially since the strong links between the hippocampal lesions and specific forms of memory were established (Scoville and Milner, 1957). This quest has spanned several scales of analysis, with efforts to link specific genes, receptors, channels and forms of cellular plasticity to learning and memory. Several studies have assessed the link between specific behavioral tasks and cellular/molecular substrates through targeted pharmacological blockades or genetic manipulations. The existence of divergent and numerous cellular/molecular components that impair *specific* learning tasks have been unveiled by these efforts, revealing considerable complexity in the plasticity networks and systems biology of learning and memory. As is evident from this complexity and associated animal-to-animal and cell-to-cell variability, which involves the ensemble of mechanisms and interactions discussed above not just from within the hippocampus but also from other brain regions, demonstrating causality with reference to learning and memory and *any one specific form of plasticity or cellular/molecular substrate*, has proven extremely challenging (Andersen *et al*., 2006; Bennett and Hacker, 2003; Bhalla, 2014; Bhalla and Iyengar, 1999; Bliss and Collingridge, 1993; Collingridge and Bliss, 1987; Jazayeri and Afraz, 2017; Kandel, 2001; Kandel *et al*., 2014; Kim and Linden, 2007; Kotaleski and Blackwell, 2010; Krakauer *et al*., 2017; Lynch, 2004; Manninen *et al*., 2010; Martin *et al*., 2000; Martinez and Derrick, 1996; Mayford *et al*., 2012; Mozzachiodi and Byrne, 2010; Neves *et al*., 2008a; Zhang and Linden, 2003).

The complexities of the networks that are involved in learning and memory are only compounded by the many-to-many mappings that are observed between behavioral observations and molecular/cellular components, the joint occurrence of several forms of plasticity with the *same* protocols (Sec. 3.6), the concurrent impairments in different forms of plasticity by blockade of the *same* signaling cascades (Sec. 3.6), the dissociations between different learning tasks and the compensatory mechanisms that are associated with the knockout of specific genes (Bailey *et al*., 2006; Jazayeri and Afraz, 2017; Krakauer *et al*., 2017; Mayford *et al*., 2012; Tsokas *et al*., 2016). For instance, the knock out of GluA1 (also referred to as GluR1 or GluRA), an AMPAR subunit that is important for expression of certain forms of synaptic plasticity, impaired only some forms of synaptic plasticity and not others at the cellular scale of analysis (Hoffman *et al*., 2002; Phillips *et al*., 2008; Zamanillo *et al*., 1999). Similarly, at the behavioral level, although behavioral deficits were observed in certain learning tasks in GluA1 knockout mice, the knock out did not alter behavior in other learning tasks (Reisel *et al*., 2002; Zamanillo *et al*., 1999). Several examples of such dissociations are reviewed in (Mayford *et al*., 2012), further emphasizing the difficulty in assigning a causal link between learning and memory and *any one specific form of plasticity or cellular/molecular substrate*.

Although this parametric and interactional complexity might seem exasperating if the goal is to pinpoint *the* cellular/molecular component that is involved in hippocampal-dependent learning and memory, it is an extremely useful substrate for the effective expression of degeneracy in achieving the goal of robust learning and memory. The ability to achieve very similar learning indices through multiple routes involving disparate forms of plasticity in several constitutive components tremendously increases the ability of the system to achieve robust learning. As a consequence of the several forms of variability and state-dependence exhibited by the learning system, in terms of the underlying components, their plasticity and combinatorial interactions, it is possible that some of these disparate routes may not involve specific cellular/molecular components or forms of plasticity in the process of achieving certain learning goals. This also implies animal-to-animal and trial-to-trial variability in the mechanisms that mediate learning, thereby calling for utmost caution in assigning one-to-one relationships between behavioral learning and specific forms of plasticity in any single brain region (Bailey *et al*., 2006; Bennett and Hacker, 2003; Jazayeri and Afraz, 2017; Krakauer *et al*., 2017; Mayford *et al*., 2012; O’Leary and Marder, 2014; Sieling *et al*., 2014; Tsokas *et al*., 2016; Vogelstein *et al*., 2014).

## 4. The causality conundrum

It is clear from the analyses above that theoretical and experimental evidence exist for: (a) several disparate combinations of distinct constitutive components elicit analogous function; (b) there are forms of animal-to-animal (channel-to-channel, neuron-to-neuron, network-to-network, etc.) variability in terms of the contributions of specific constitutive components that mediate similar function; and (c) the components that mediate similar function, and their relative contributions to the emergence of this function are state-dependent, and could undergo experience-dependent plasticity (towards maintaining robustness of that function or towards learning-dependent alteration of function). Juxtaposed against these observations is the question on whether it is even possible to exclusively assign causal one-to-one relationships between function and specific constitutive components. Evidence for the existence of degeneracy, variability and adaptability have made us acutely aware of the possibility that we could be committing mereological fallacies (Bennett and Hacker, 2003; Varzi, 2016), whereby we assign specific behavioral roles to parts of the animal’s brain or to plasticity therein (Bailey *et al*., 2006; Jazayeri and Afraz, 2017; Krakauer *et al*., 2017; Mayford *et al*., 2012; O’Leary and Marder, 2014; Sieling *et al*., 2014; Tsokas *et al*., 2016; Vogelstein *et al*., 2014).

### 4.1. Inevitable flaws in an experimental plan to establish causality that leaps across multiple scales

Let us chart a hypothetical experimental plan where we are interested in demonstrating that a specific form of learning behavior is dependent on plasticity in one specific component (let’s say component X) in a brain region of our choice (let’s say hippocampus). We first measure *in vivo* plasticity in component X along with its time course, and let us say that we find a prominent correlation between this time course and the time course of behavioral learning. Next, we introduce an established blocker of plasticity in component X specifically into the hippocampus, and find that this blocks both the plasticity in component X *in vivo* and impairs learning. We repeat similar experiments with (a) an established pharmacological blocker of component X infused into the hippocampus; (b) transgenic manipulations that take out component X completely in the hippocampus; (c) a pharmacological blocker that leaves component X intact, but impairs its plasticity by blocking a mechanism that induces plasticity in component X; and (d) genetic knockout of mechanisms that mediates plasticity in component X. Let’s say that learning was impaired in all four cases, and there was no plasticity in component X in the last two cases (in the first two cases component X was completely abolished). As a final nail in the hypothesis to link plasticity in hippocampal component X to the specific learning behavior, we artificially alter component X and consequently find behavioral signatures related to the learning process. Therefore, we have shown that component X and its plasticity are necessary and sufficient for the specific learning behavior. This experimental plan is broadly similar to that proposed by (Stevens, 1998) to test the hypothesis that auditory synapses in the amygdala become strengthened by LTP during behavioral training that attaches “fear” to the tone, and that he memory of the tone as a fear-producing stimulus resides in the strength of the synapses from the auditory thalamus (Stevens, 1998):

> “How could this idea be tested? It should be that (1) blocking LTP prevents fear learning; (2) the sensory pathways from the thalamus and cortex to the amygdala are capable of LTP; (3) auditory fear conditioning increases the amygdala’s postsynaptic response to the tone, and these increases are prevented by blocking LTP pharmacologically or in another way; and (4) inducing LTP in the thalamoamygdaloid pathway attaches “fear” to appropriate sensory stimuli.”

Although this experimental plan has shown that component X and its plasticity are necessary and sufficient for the specific learning behavior, given the complexity that we have elucidated thus far, this experimental design *does not* provide a causal link between component X or its plasticity with behavior. First, we were so focused on component X that we implicitly precluded the change of any other component either in the hippocampus or in other brain region. Given the rich complexity in the distinct components, their plasticity and interactions among them, it is infeasible that only component X in the hippocampus was changing in response to the behavioral stimulus. It is now well established that several cellular components change in response the same calcium signal or the activation of the same signaling cascade, and there are several parallel homeostasis mechanisms that also exhibit degeneracy. This implies that altering component X in the hippocampus *without* altering anything else across the brain is highly unlikely. Therefore, if we had performed the same set of experiments on another component Y, we might have arrived at similar conclusions (including correlated time courses). In other words, it is important not to interpret measurement correlations as evidence for causation, and to understand that absence of measurements in other forms of plasticity or plasticity in other brain regions does not mean they don’t coexist with the form of plasticity that we are focused on.

Second, when we blocked plasticity in component X, given the complexities elucidated above, it is highly unlikely that we *specifically* blocked plasticity in component X without disturbing plasticity in any other measurement or without introducing metaplasticity in some other form of plasticity (Sec. 3.6–3.7). For instance, from a cellular perspective, theta burst pairing results in plasticity of synaptic strength and of HCN, *A*-type K^+^ and SK channels, and pharmacologically blocking NMDAR receptors impairs plasticity not just in one of them, but in *all* of them (Fan *et al*., 2005; Frick *et al*., 2004; Lin *et al*., 2008; Losonczy *et al*., 2008). Thus if we had observed impairment of plasticity in only one of these components, we would have wrongly concluded that to be the only component that changes with TBP. Returning to our experimental plan on the role of component X, there could several other secondary and unintended effects of blocking plasticity in component X that spans the hippocampus and other brain regions (Bhalla, 2014; Jazayeri and Afraz, 2017; Kotaleski and Blackwell, 2010; Krakauer *et al*., 2017; Otchy *et al*., 2015). Thus, it is prudent not to dismiss absence of measurements as absence of evidence for other components.

Third, when we performed the experiment of artificially altering component X, it is obvious that it is highly unlikely that we achieved this without disturbing any other component in some brain region or without introducing metaplasticity in some form of plasticity. Therefore, the alternate interpretations of our observations (other than the “linear narrative” that concludes “plasticity in hippocampal component X mediates learning behavior”) are innumerable given the staggering complexity of the underlying system and the degeneracy involved in accomplishing the learning task. Ruling out *all these* alternate interpretations is essential for convergence to the linear narrative, but is rather impossible because measurements of all constitutive components in all brain regions is currently infeasible. From a nonlinear dynamical system perspective (Guckenheimer and Holmes, 1983; Nayfeh and Balachandran, 1995; Strogatz, 2014), our “linear narrative” and the associated inference are equivalent to declaring a component to be critically important for system performance because perturbation to that one component, — which is part of a high-dimensional, adaptive, non-linear dynamical system with strong coupling across dimensions, — collapses the system. Additionally, especially given the expression of degeneracy, in our artificial perturbation experiment, we showed that the system *could* perform a specific behavior when we introduced a perturbation to component X. However, this observation does not necessarily imply that the system *does* employ a similar perturbation to component X to elicit the same behavior under normal ethological conditions (Adamantidis *et al*., 2015). Given the degeneracy framework, it is important to appreciate that the existence of *a* solution neither implies its uniqueness nor does it ensure that the solution is employed by the physiological system under standard ethological conditions.

### 4.2. Degeneracy: The way forward

It is important to distinguish between understanding functionality that emerges through interactions between components in an adjacent scale and efforts aimed at causality that leaps across multiple scales. It is clear that assessing interactions between constitutive components in the emergence of function in an adjacent scale have provided invaluable insights in neuroscience. As an example, the question on how different ionic currents at the molecular scale interact to result in the emergence of an action potential in the cellular scale (Hodgkin and Huxley, 1952) has revolutionized several aspects of neuroscience over the past several decades. Even within the framework of degeneracy, the question on whether and how different combinations of disparate combinations of parameters in a give scale could result in similar functionality in an adjacent scale have provided deep insights into how the nervous system might be solving the robustness problem in the face of variability (Anirudhan and Narayanan, 2015; Dhawale *et al*., 2017; Foster *et al*., 1993; Gjorgjieva *et al*., 2016; Goldman *et al*., 2001; Katz, 2016; Marder, 2011; Marder and Goaillard, 2006; Marder *et al*., 2015; Marder and Taylor, 2011; Mukunda and Narayanan, 2017; O’Leary and Marder, 2014; Prinz *et al*., 2004; Rathour and Narayanan, 2012a, 2014; Taylor *et al*., 2009).

However, causal leaps beyond a single scale of analysis should be treated with extreme caution. For instance, approaches assuming a unique reductionist solution for a behavioral observation will invariably end up in apparently contradictory conclusions about *the* mechanism that mediates behavior. Prominent among the several reasons that result in these apparent contradictions — such as adaptive compensations and animal-to-animal variability — is inherent degeneracy, where disparate combinations of components could result in identical behavior in a manner that is dependent on several factors, including behavioral state. The flaws that emerge in an experimental plan to establish causality that leaps multiple scales in a nonlinear dynamical system that expresses degeneracy are obvious from the analysis presented above. Here, it is critical to ask the impossible question on whether we are sure that *nothing else* has changed in neurons (and other cells) of the same brain region or the other, which could be mediating/contributing to the observed behavioral changes *before* declaring a causal one-to-one relationship between a molecular/cellular component and behavior.

This is especially important because there are several properties that *emerge* at each jump along the multiscale axis of neuroscience (Fig. 1A), and leaps across multiple scales (like genes to behavior) traverses several *emergent* properties owing to innumerable nonlinear processes that exhibit degeneracy. This yields a system that is intractable even at the scale where the perturbations were introduced because of the complex feedback loops spanning several scales that mediate homeostasis and adaptation. Consequently, the outcomes of any perturbation at any scale are critically dependent on several components across scales, the nature of interactions of these components with the perturbation and importantly on the adaptation that is triggered by the perturbation in all these components across scales. Therefore, extreme caution should be exercised in assigning causal one-to-one relationship between components (or manifolds) that are several scales apart along the multi-scale axis (Bennett and Hacker, 2003; Jazayeri and Afraz, 2017; Krakauer *et al*., 2017; Otchy *et al*., 2015).

Together, while degeneracy is an invaluable asset to evolution, physiology and behavior in achieving robust functions through several degrees of freedom, it makes the resultant complex system rather intractable. This intractability makes it nearly impossible to achieve the goals of reductionism, where the pursuit has largely been for causal one-to-one relationships that leap across several scales. Several thorny debates in the field about apparent contradictions involving different components mediating the same function could be put to rest if this requirement of one-to-one relationships is relaxed. Specifically, the ubiquitous expression of degeneracy spanning multiple scales offers an ideal reconciliation to these controversies, through the recognition that the distinct routes to achieve a functional goal are not necessarily contradictory to each other, but are alternate routes that the system might recruit towards accomplishment of the goal. The intense drive to make leaps across multiple scales to establish unique one-to-one relationships should instead be replaced by a steadfast recognition for degeneracy as an essential component in physiology, behavior and evolution. This recognition, apart from precluding one-to-one relationships, would provide clear warnings in assigning causal relationships that leap across multiple scales and multiple emergent properties. Importantly, this recognition would pave the way for a strong focus on integrative and holistic treatises to neuroscience and behavior, arguments for which have only been growing over the years (Bennett and Hacker, 2003; Edelman and Gally, 2001; Jazayeri and Afraz, 2017; Krakauer *et al*., 2017; Tononi and Edelman, 1998; Tononi *et al*., 1998; Tononi *et al*., 1994; Tytell *et al*., 2011). Future approaches should recognize that behavior emerges from disparate combinations of tightly cross-coupled multi-scale emergent properties, each diverging and converging at each scale of analysis through degeneracy spanning complex parametric and interactional spaces. Large-scale databases related to neuronal morphology, models and physiology — such as the Allen brain atlas (Sunkin *et al*., 2013), ICGenealogy (Podlaski *et al*., 2017), Channelpedia (Ranjan *et al*., 2011), Neuromorpho (Ascoli *et al*., 2007), ModelDB (Hines *et al*., 2004) and Neuroelectro (Tripathy *et al*., 2014) — provide ideal tools for such analyses involving large parametric spaces, and could provide critical insights about the role of degeneracy in the emergence of robust brain physiology and its links to behavior.

## 5. Conclusions

In this review, we systematically presented lines of evidence for the ubiquitous expression of degeneracy spanning several scales of the mammalian hippocampus. We argued that the framework of degeneracy in an encoding system shouldn’t be viewed from the limited perspective of maintaining homeostasis, but should be assessed from the perspective of achieving the twin goals of encoding information and maintaining homeostasis. Within the broad framework of degeneracy, it is extremely important that future studies focus on the fundamental questions on (i) how does the brain change its constituent components towards encoding new information without jeopardizing homeostasis?; and (ii) how do homeostatic mechanisms maintain robust function without affecting learning-induced changes in the brain? Without an effective answer to this overall question on concomitant learning and homeostasis in the face of staggeringly combinatorial complexity, our understanding of the nervous system in terms of its ability to systematically adapt to the environment will remain incomplete. Although the core conclusions on degeneracy reviewed and analyzed here would extend to other mammalian brain regions and functions that they have been implicated in encoding processes, this extrapolation should be preceded by careful assessment of the specifics associated with the constitutive components and specific interactions there. Additionally, although our focus here was on encoding, homeostasis and physiology, it is important that future studies also assess the implications for degeneracy in the emergence of pathological conditions (Edelman and Gally, 2001; O’Leary *et al*., 2014).

Finally, returning to the distinction between the “structure defines function” and the “form follows function” perspectives, it seems like the distinction also seemingly extends to the methodology that is deemed appropriate for assessing neuronal systems. At one end, a strong emphasis is placed on the requirement for an experimental approach (Buzsaki, 2006):

> “The complexity and precision of brain wiring make an experimental approach absolutely necessary. No amount of introspection or algorithmic modeling can help without parallel empirical exploration.”

At the other end, the emphasis, reflecting Richard Feymann’s quote “What I cannot create, I do not understand”, is on *in silico* approaches (Sakmann, 2017):

> “At present however, it seems that “What we cannot reconstruct *in silico* and model we have not understood”.”

Within the degeneracy framework, however, it is starkly evident from existing literature reviewed here that a holistic combination of computational and experimental techniques is indispensible towards understanding structure-function relationships and the associated complexities (Das *et al*., 2017; Edelman and Gally, 2001; Foster *et al*., 1993; Marder, 1998, 2011; Marder and Goaillard, 2006; Marder and Taylor, 2011; Rathour *et al*., 2016; Rathour and Narayanan, 2012a, b, 2014; Sporns *et al*., 2000; Tononi and Edelman, 1998; Tononi *et al*., 1998; Tononi *et al*., 1994, 1996, 1999).

Emphasizing the strong links between biology and evolution, Theodosius Dobzhansky had written “nothing in biology makes sense except in the light of evolution” (Dobzhansky, 1973). Given the ubiquitous prevalence of degeneracy and its strong links to evolution (Edelman and Gally, 2001), it is perhaps apt to add a corollary to this quote and state “nothing in physiology makes sense except in the light of degeneracy”.

## ACKNOWLEDGMENTS

We thank Dr. Peter Jedlicka, Dr. Manisha Sinha and members of the cellular neurophysiology laboratory for useful discussions and for thoughtful comments on a draft of this manuscript. Work reviewed here was supported by the Wellcome Trust-DBT India Alliance (Senior fellowship to RN: IA/S/16/2/502727), Human Frontier Science Program (HFSP) Organization (RN), the Department of Biotechnology (RN), the Department of Science and Technology (RN) and the Ministry of Human Resource Development, India (RKR and RN).

